# Transcription activator-coactivator specificity is mediated by a large and dynamic fuzzy protein-protein complex

**DOI:** 10.1101/221747

**Authors:** Lisa M. Tuttle, Derek Pacheco, Linda Warfield, Jie Luo, Jeff Ranish, Steven Hahn, Rachel E. Klevit

## Abstract

Transcription activation domains (ADs) are inherently disordered proteins that often target multiple coactivator complexes, but the specificity of these interactions is not understood. Efficient activation by yeast Gcn4 requires tandem Gcn4 ADs and four activator-binding domains (ABDs) on its target, the Mediator subunit Med15. Multiple ABDs are a common feature of coactivator complexes. We find that the large Gcn4-Med15 complex is heterogeneous, containing nearly all possible AD-ABD interactions. This complex forms using a dynamic fuzzy protein-protein interface where ADs use hydrophobic residues to bind hydrophobic surfaces of the ABDs in multiple orientations. This combinatorial mechanism allows individual interactions of low affinity and specificity to generate a biologically functional, specific, and higher affinity complex despite lacking a defined protein-protein interface. This binding strategy is likely representative of many activators that target multiple coactivators and allows great flexibility in combinations of activators that synergize to regulate genes with variable coactivator requirements.

## INTRODUCTION

Transcription activators lie at the endpoint of signaling pathways that control transcription in response to development, cell growth, stress, and other physiological signals (Levine et al., 2014; Spitz and Furlong, 2012). A central challenge in understanding gene regulation is to determine the mechanism and specificity of activators. Transcription activation domains (ADs) are intrinsically disordered, lack a stable structure in the absence of a binding partner, and do not share obvious primary sequence similarity (Dyson and Wright, 2005; Hahn and Young, 2011; Nguyen Ba et al., 2012; Tantos et al., 2012; Tompa et al., 2014). Most known activators work by targeting transcription coactivator complexes to stimulate both transcription preinitiation complex (PIC) assembly and chromatin modifications (Hahn and Young, 2011; Weake and Workman, 2010). For over thirty years, a central mystery surrounding transcriptional activators, once referred to as “acidic blobs and negative noodles” (Sigler, 1988), is how they specifically target unrelated coactivators?

The acidic transcription activator Gcn4 from *S. cerevisiae* has properties common to many eukaryotic activators. Gcn4 contains tandem intrinsically disordered ADs (tADs) that target different coactivators including Mediator, SAGA, Swi/Snf and/or NuA4 to regulate a large number of genes in response to metabolic stress (Brown et al., 2001a; Brzovic et al., 2011; Drysdale et al., 1995; Fishburn et al., 2005; Herbig et al., 2010; Jackson et al., 1996; Jedidi et al., 2010; Natarajan et al., 2001; Qiu et al., 2016; Swanson et al., 2003; Warfield et al., 2014; Yoon et al., 2003). A key Gcn4 target is Med15/Gal11, a component of the Mediator tail module (Brzovic et al., 2011; Herbig et al., 2010; Jedidi et al., 2010; Warfield et al., 2014). Within its N-terminal half, Med15 contains four structured domains: a KIX domain (Novatchkova and Eisenhaber, 2004; Yang et al., 2006) and activator-binding domains (ABDs) 1, 2, and 3 that are recognized by Gcn4 (Herbig et al., 2010; Jedidi et al., 2010) (**Fig. 1A**). Although interactions have been detected between single AD sequences and individual ABDs, no single AD-ABD interaction is sufficient for efficient stimulation of transcription; high levels of activation are achieved only with the complete Med15 activator binding region and the tandem Gcn4 ADs (Herbig et al., 2010; Jedidi et al., 2010). This implies that a single AD-ABD interaction lacks sufficient affinity and/or specificity to promote transcription.

**Figure 1.**
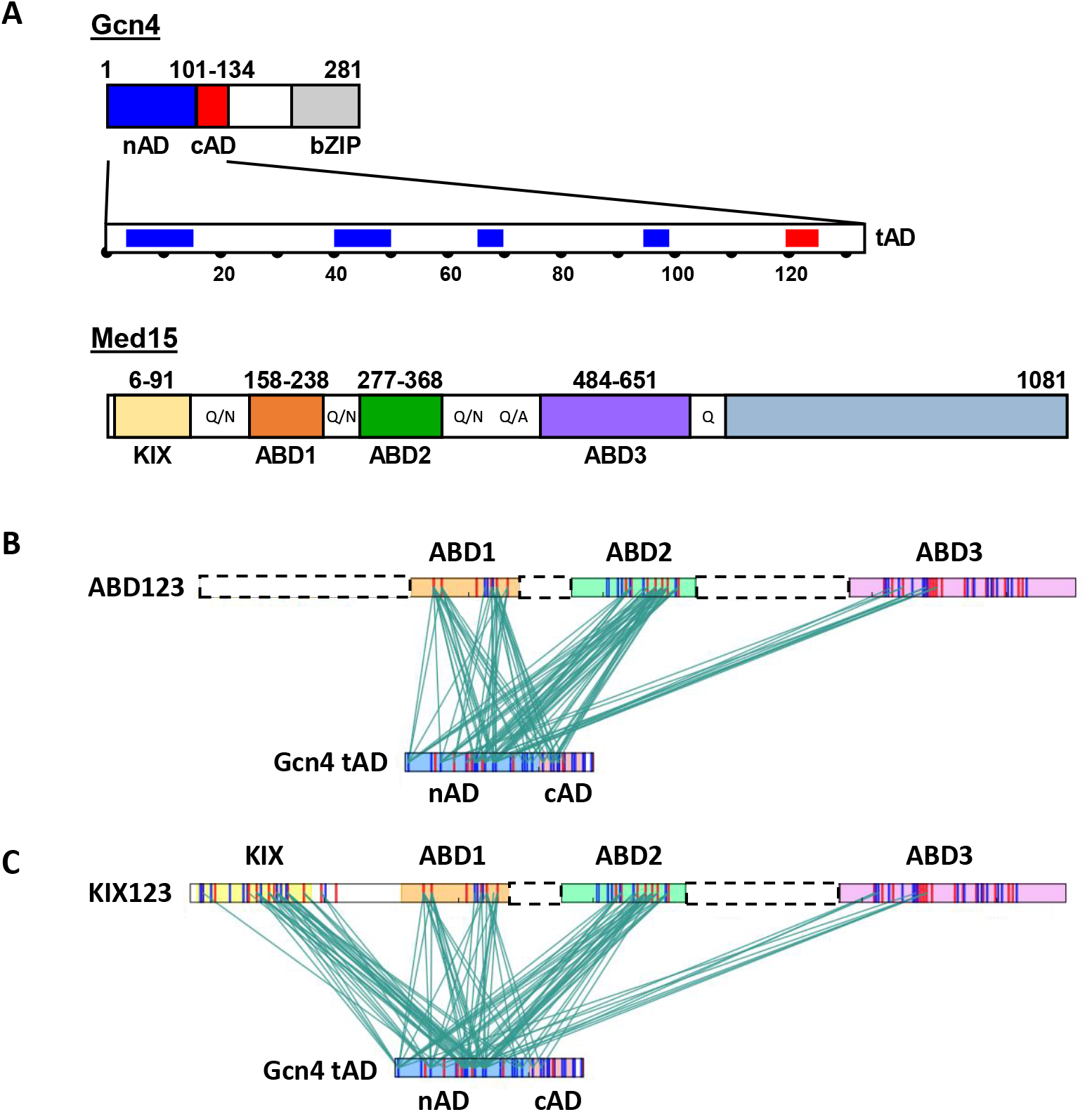
Gcn4 tAD crosslinks to each Med15 ABD and KIX. **(A)** Gcn4 consists of the tAD and bZIP subdomains. The expanded view of the tAD highlights key hydrophobic residue clusters located in the nAD (blue) and cAD (red). Med15 contains four structured domains in the activator-binding region and the long Q-rich linkers between the domains are indicated. **(B)** ABD123-tAD and (C) KIX123-tAD crosslinking results. Lines between the Med15 and Gcn4 constructs indicate the sites of crosslinking that were identified by mass spectrometry. Dashed boxes indicate deleted regions of the wild-type sequence for each Med15 construct. Red and blue bars within each subdomain indicate acidic and lysine residues, respectively.

The use of multiple ADs and ABDs is not unique to Gcn4 and Med15. Other strong activators such as VP16 and p53 contain tandem ADs and coactivators such as CBP, Swi/Snf, and SAGA/NuA4 all contain multiple ABDs (Brown et al., 2001b; Chang et al., 1995; Diaz-Santin et al., 2017; Ferreon et al., 2009; Langlois et al., 2008; Prochasson et al., 2005; Regier et al., 1993; Unger et al., 1992; Walker et al., 1993). While multiple ADs and ABDs appears to be a common theme in transcriptional regulation, the mechanism of how a multiplicity of AD-ABD interactions give rise to the affinity and specificity required for AD function remains undefined.

We previously showed that the Gcn4 central AD (cAD) binds to ABD1 with a fuzzy protein-protein interface. Disordered on its own, the cAD peptide binds in multiple orientations to a shallow hydrophobic groove on ABD1, mediated almost entirely by hydrophobic interactions (Brzovic et al., 2011). Because a single AD-ABD interaction is not sufficient for transcription activation (Herbig et al., 2010), an important question arising from our findings is whether a fuzzy binding mode is retained in the larger physiological Gcn4-Med15 complex with its full complement of ADs and ABDs. An alternative model predicts that complexes with multiple ADs and ABDs lock into a specific, more stable complex with higher affinity binding and conventional protein-protein interfaces. Indeed, in two systems involving intrinsically disordered proteins (IDPs) and structured binding partners that have been characterized at a structural level, the IDPs form heterogeneous complexes with individual interacting partners, but form stable distinct higher-ordered structures when all binding partners are present (Dyson and Wright, 2016; Saio et al., 2014). Such examples illustrate the importance of examining large natural complexes of IDPs and their targets. The distinction between the two possible mechanisms has important implications for which AD sequences can function together and the specificity of AD targets. For example, a fuzzy binding mechanism is predicted to allow greater flexibility in AD-AD and AD-ABD combinations compared to systems that lock into a specific complex.

To determine whether the Gcn4-Med15 complex forms a specific higher-order complex or interacts via a large fuzzy interface, we characterized the interactions between the Gcn4 tandem ADs (tADs) and the four ABDs of Med15, both individually and as part of the larger Gcn4-Med15 complex. Our data demonstrate that the presence of multiple binding components leads to a large gain in the affinity of Gcn4-Med15 binding, but does not lead to formation of a distinct higher-order structure. The functional complex is a dynamic fuzzy “free-for-all” involving the hydrophobic patches of the tADs with ABD 1–3 and the KIX domain. Our results provide a general model for the mechanism of many activators that functionally bind coactivators with multiple ABDs and demonstrate how multiple weak fuzzy interactions synergize to generate a biologically functional and specific interaction without adopting a unique protein-protein interface.

## RESULTS

### Both Gcn4 ADs interact with all individual Med15 ABDs

As a first step in characterization of the complete Gcn4-Med15 complex, we examined regions required for AD function and investigated the affinity of the individual and higher order AD-ABD interactions. Gcn4 contains two ADs, termed the N-terminal AD (nAD; residues 1–100) and central AD (cAD; residues 101–134) (Drysdale et al., 1995) (**Fig. 1A**). The cAD is one of the shortest ADs known with most of its function contained within a five-residue sequence motif WxxLF (residues 120–124) (Warfield et al., 2014). In contrast, nAD function is dependent on clusters of hydrophobic residues located throughout residues 1–100 (Jackson et al., 1996). As AD function is usually confined to much shorter sequences, we examined whether any of the individual hydrophobic regions within the nAD have function and whether any non-hydrophobic regions of the nAD contribute to activity. A series of deletions and alanine substitutions was made within a Gcn4 derivative lacking the cAD and function was measured at *ARG3* and *HIS4*, two Gcn4-dependent yeast genes (**Fig. S1**). Consistent with prior results (Jackson et al., 1996), our analysis showed that Gcn4 nAD function requires four short hydrophobic regions (residues 4–16, 40–49, 65–69 and 94–98). We also found that residues between the clusters can be deleted with little or no detrimental effect and, in several cases, the deletions increase activity. In particular, deletion of residues between the first three hydrophobic regions to create one long hydrophobic region generates an AD with up to 4.6-fold higher activity relative to wild type. Importantly, none of the individual short hydrophobic regions has substantial activity when fused to the Gcn4 DNA binding domain. Therefore, nAD function arises from multiple short non-redundant hydrophobic clusters with little or no inherent activity that together generate AD function.

We used isothermal calorimetry (ITC) and/or fluorescence polarization (FP) to measure the binding affinity of each AD and individual ABD, as well as the binding affinity of a tAD-ABD1,2,3 complex (Table 1). With one exception, all the individual pairwise interactions between individual nAD and cAD peptides and individual Med15 ABDs have similar low/modest affinities (K_d_ values of ∼3–20 μM). The exception is cAD binding to ABD2, which is ∼10-fold weaker (150 μM K_d_). Although the Med15 KIX domain is as important functionally for Gcn4 activation as any other ABD (Herbig et al., 2010), neither nAD or cAD showed detectable KIX binding in our assays. The tandem AD polypeptide (Gcn4 residues 1–134) binds to each individual ABD with an affinity close to the stronger of the individual nAD or cAD interactions, suggesting that the individual Gcn4 ADs bind to the same or overlapping sites on the ABDs. A striking gain in affinity is observed when the tAD binds to longer Med15 polypeptides containing multiple ABDs (ABD123 and KIX123). For example, tAD binding to ABD123 (ABD1, ABD2, and ABD3 connected by short linkers) is about 20-fold higher affinity than the strongest piecemeal interaction, signifying that, when all domains are present, more than one ABD simultaneously contributes to binding. Notably, the presence of KIX along with ABD1,2,3 (KIX123) increases Gcn4 affinity by ~30%. Altogether the results show that multiple weak AD-ABD and KIX interactions in the full-length proteins combine to yield a much higher affinity Gcn4-Med15 interaction – in agreement with in vivo activation function.

**Table 1.**
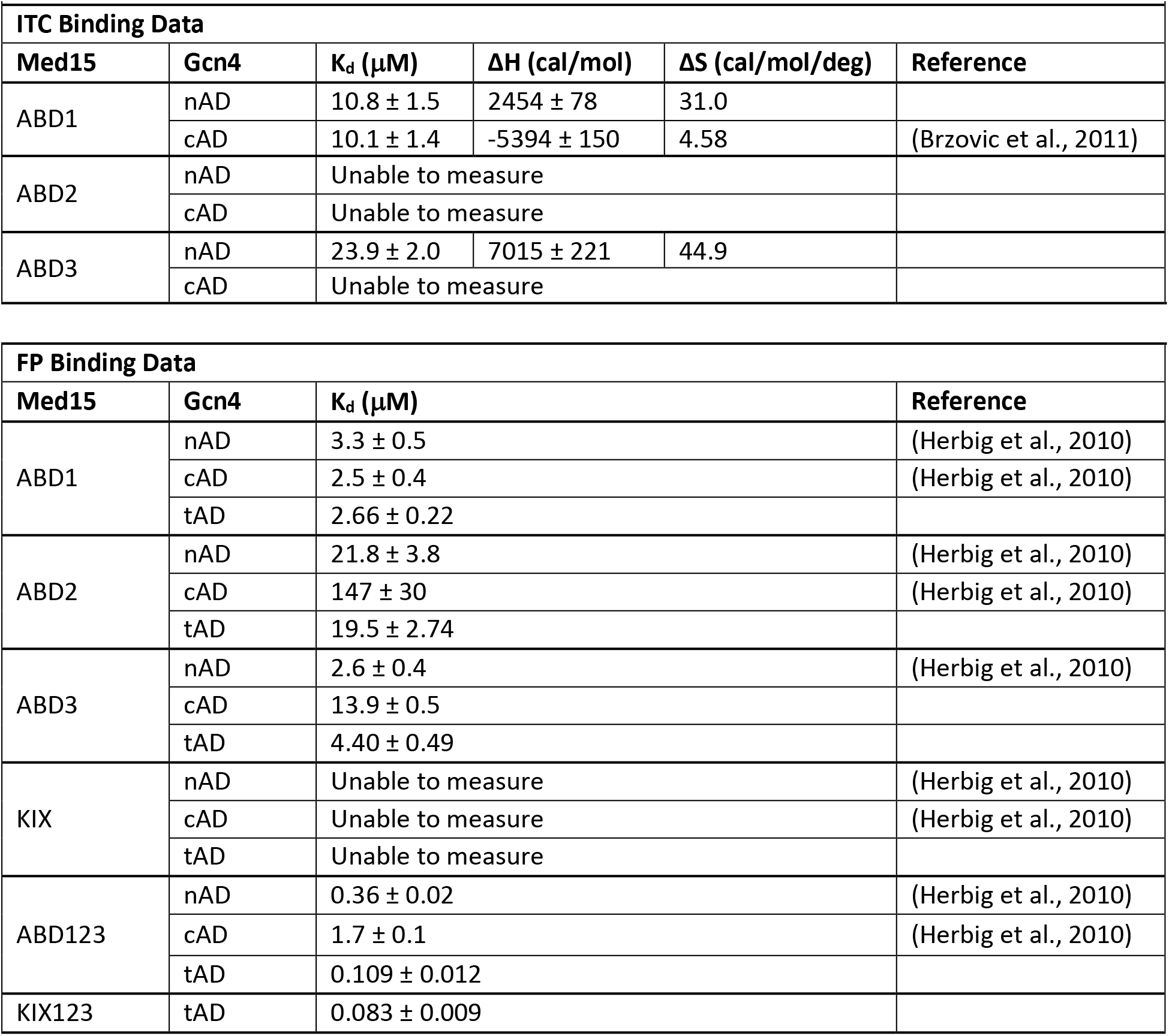
Affinity of Gcn4 ADs for Med15 activator-binding domains

### A heterogeneous Gcn4-Med15 complex

A prediction of the fuzzy complex model for Gcn4-Med15 is that a complex containing multiple ADs and ABDs will not contain fixed protein-protein interactions and that individual ADs will sample multiple ABDs in different orientations as was observed for cAD-ABD1 (Brzovic et al., 2011). The alternative “lock-down” model predicts that specific protein-protein interactions will be adopted in the larger complexes. To test the two models, we used chemical crosslinking analyzed by mass spectrometry to detect interactions between the Gcn4 tAD and Med15 ABD123 or KIX123 (**Fig. 1B, C**). Complexes of Gcn4 residues 1–140 (tAD) with either of the two Med15 constructs were treated with the zero-length crosslinker EDC that links amine groups, such as those found in lysine side chains, to carboxylate groups, such as those found in the side chains of acidic amino acids. Identified crosslinks are summarized in **Figure 1B,C** and reported in **Tables S1 and S2**.

Crosslinks were observed between both ADs of Gcn4 and all four defined structural regions of Med15 (KIX domain and ABD1, 2, 3). Strikingly, pairwise linkages are detected between all possible domains with one exception: no crosslinks between Gcn4 cAD and Med15 ABD3 were detected. In general, fewer crosslinks were identified involving ABD3 than those involving the other subdomains, even though ABD3 has a stronger intrinsic affinity for AD than does ABD2 (**Table 1**) and has a similar number of potential crosslinking sites. The KIX domain, with little or no detectable binding to Gcn4 on its own, crosslinks extensively to Gcn4 in the large complex, mainly to nAD. Notably, the crosslink patterns between ABD1–3 and AD regions are unchanged when the KIX domain is also present (KIX123 compared to ABD123). Thus, the crosslinking data show that KIX interacts directly with Gcn4, consistent with functional assays showing that KIX is as important as the individual ABD1, 2, and 3 domains for Gcn4 activation (Herbig et al., 2010). The results reveal that interactions involving the weaker binding domains of Med15 occur even when all domains are present, i.e., no domain outcompetes the others for AD binding. Such a mechanism predicts transient binding for each AD-ABD/KIX interaction.

### tAD-Med15 interactions are a sum of the individual nAD and cAD interactions

Together, the crosslinking and binding assays indicate that while the large Gcn4-Med15 complex has high affinity, it is nevertheless heterogeneous. Two models are consistent with this behavior: (1) each AD binds each ABD/KIX with a dynamic and fuzzy protein-protein interface, or (2) AD-ABD binding is dynamic, but individual AD-ABD binding occurs with unique non-fuzzy interfaces. To distinguish between these models, we investigated the properties of the individual AD-ABD complexes by NMR and compared the results to those of the complete complex.

We first examined the environment of the polypeptide backbones of the nAD, cAD, and tAD upon Med15 binding. Titrations were carried out for each combination of ^13^C,^15^N-labeled nAD, cAD, and tAD binding to unlabeled ABD1, ABD2, ABD3, and ABD123. In these experiments, effects on the isotopically-labeled component due to binding the non-labeled (silent) component are observed in (^1^H,^15^N)- and (^1^H,^13^C)-HSQC spectra. Titrations were taken to full saturation where possible, based on measured K_d_ values (Table 1).

In each titration series, NMR peaks for affected residues changed their resonance position continuously as a function of added binding partner until saturation was reached (e.g., **Fig. S2**). The chemical shift perturbation (CSP) of a peak was calculated as the difference in its chemical shift in the spectrum of free AD and in the most saturated spectrum (Methods). CSPs for AD backbone amide (NH) groups are displayed as histograms in **Figure 2**, with a panel for each individual ABD (panels **A-C**) and a panel for the combined ABD123 (panel **D**). Previously reported results for ^15^N-cAD:ABD1 are included here for a complete comparison (Brzovic et al., 2011). Several general features emerge from the large body of data. First, perturbations occur in hydrophobic residue clusters along the AD sequence that correspond to the previously identified hydrophobic regions in Gcn4. Second, the same clusters are perturbed regardless of which ABD has been added. Third, in all cases, the fourth nAD hydrophobic cluster (residues 94–98) shows the largest CSPs, followed by the cAD cluster (residues 120–124), with smaller shifts in each of the other hydrophobic regions. Fourth, there are no new perturbations associated with tAD binding to Med15; both the identity and magnitudes of the chemical shift perturbations (CSPs) observed in the isolated nAD and cAD are retained in full-length tAD (**Fig. 2**, left and right histograms are highly similar). Thus, we find no additional cryptic interaction sites in the full-length Gcn4 tandem AD.

**Figure 2.**
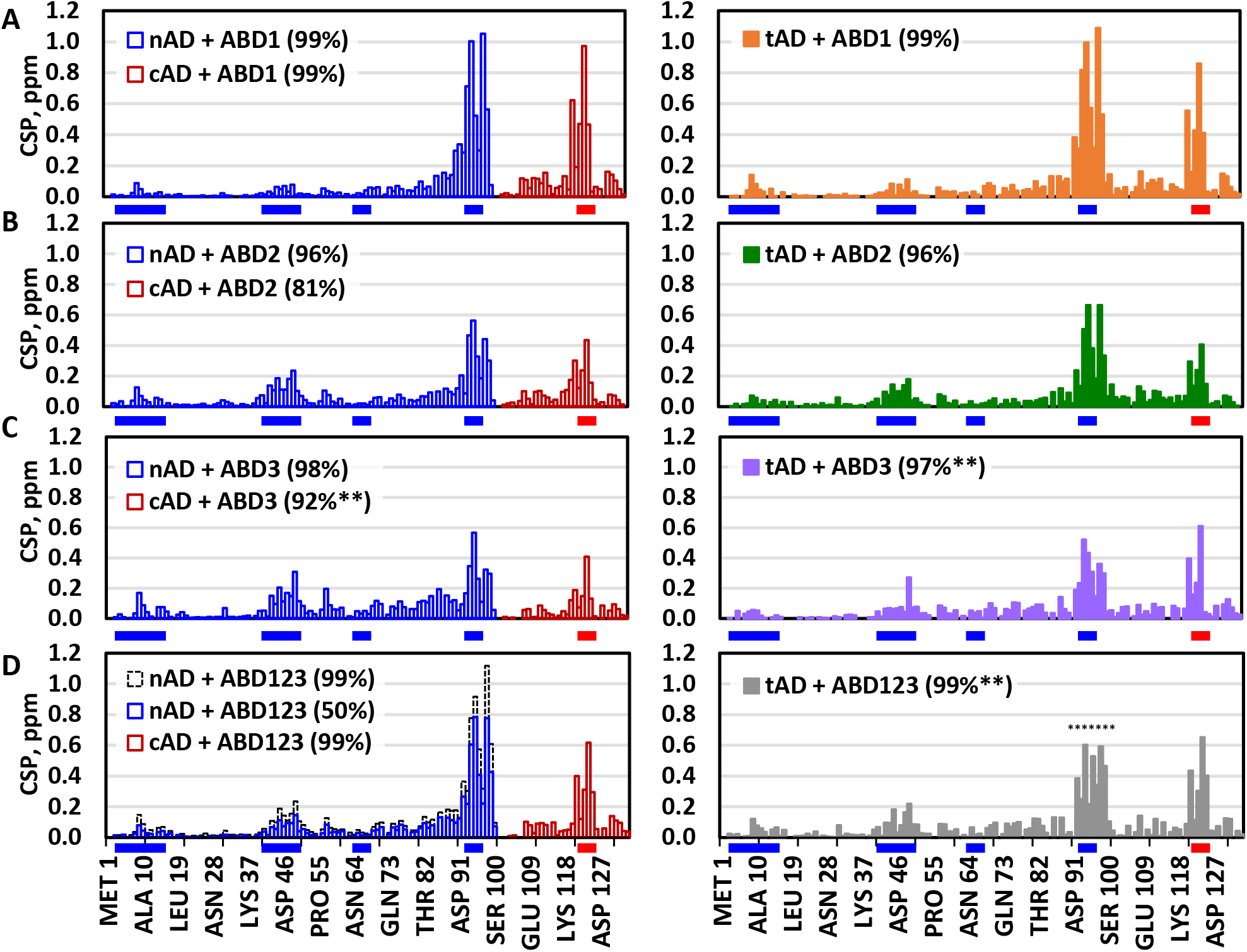
Gcn4 nAD and cAD independently interact with Med15 ABDs. **(A-D)** The NH Chemical Shift Perturbation (CSP) of Gcn4 upon addition of Med15. For each ABD, nAD and cAD regions have similar CSPs as independent regions (left) or in the context of full-length tAD 1–134 (right). See **Fig. S2** for (^1^H,^15^N)-HSQC titration spectra. **(C)** Many cAD region peaks are lost after ~50% saturation for both the cAD + ABD3 and tAD + ABD3 titrations. **(D)** HSQC peaks for tAD residues 91–99 of the tAD + ABD123 titration are lost at saturation levels > 50% so CSPs for nAD + ABD123 are also shown at 50% saturation for comparison; dashed black lines show full saturation data for nAD+ABD123 in **D**. In all panels, ** indicate situations where not all peaks trajectories could be followed to complete saturation. Hydrophobic regions are highlighted by blue (nAD) and red (cAD) bars beneath each plot.

While the same regions of the ADs show chemical shift perturbations when binding to all ABDs, the magnitudes of the shifts depend on the ABD binding partner. The contribution of each cluster is parsed out in **Figure S3A**, in which the maximum CSPs in each cluster are compared. Two clusters dominate the binding to all ABDs, one in the nAD and cAD. The other nAD clusters contribute differently depending on the ABD: the larger CSPs for the weaker hydrophobic regions in the ABD2 and ABD3 titrations suggests that these clusters play a larger role in ABD2 and ABD3 binding than in ABD1. The CSPs of the weakly interacting hydrophobic regions of nAD do not increase in the full-length complex, as might occur if a sequential binding mechanism were in play. Instead, the CSPs in tAD:ABD123 resemble the component tAD:ABD complexes, indicating that the importance of the weaker hydrophobic clusters for activator function is not a result of new or stabilized interactions in the full-length AD-ABD complex.

### Gcn4 ADs become helical upon binding Med15 ABDs

The primary interacting regions of many ADs become helical upon binding to their targets; e.g., residues 117–125 in the Gcn4 cAD adopt helical secondary structure upon ABD1 binding (Brzovic et al., 2011). We used the chemical shifts of free and bound nAD and cAD to assess secondary structure upon binding to Med15 ABDs (Tamiola and Mulder, 2012). Unbound nAD and cAD are completely disordered on their own, but each gains substantial helical content upon binding ABD1 (**Fig. S3B**). The two Gcn4 C-terminal hydrophobic clusters gain the most helicity: the nAD cluster exhibits up to 50% helical content and the cAD cluster exhibits about 30%, while each of the other three nAD hydrophobic clusters show some helical character (<20%) upon binding ABD1. These analyses were performed on the two individual ADs where the most complete resonance assignments are available. However, chemical shifts in the tAD essentially match the component shifts of nAD and cAD (**Fig. 2**), allowing us to conclude that the same pattern and magnitude of helical structure is gained when tAD binds to ABD1.

Secondary structure gained by nAD upon binding to ABD2 was also calculated and qualitatively resembles that of nAD-ABD1, except that the C-terminal hydrophobic cluster shows only ~25% helical content (**Fig. S3B**). As this analysis was performed for spectra collected under saturating concentrations of each ABD, the differences in estimated helicity of residues 94–98 suggest either that this cluster spends less time bound to ABD2 than it does to ABD1 (i.e., has a faster off-rate) and/or that it binds to ABD2 in non-helical as well as helical conformations. Overall, our analysis shows that: (1) hydrophobic regions of ADs that interact with ABDs gain helical content upon binding and (2) the most frequent AD-ABD contacts are with Gcn4 hydrophobic clusters 94–98 and 120–124. Nevertheless, the other nAD hydrophobic clusters play an essential functional role (**Fig. S1**) and contribute significantly to the overall interaction with Med15.

### Structure of Med15 ABD2

To determine the Gcn4-Med15 binding mechanism, CSPs in the ABDs upon Gcn4 binding were determined. Unlike the ADs which are intrinsically disordered in their free states, ABDs typically adopt defined three-dimensional structures. However, few ABD structures have been experimentally determined, making structure prediction unreliable. We next set out to determine structures for Med15 ABD2 (residues 277–368) and ABD3 (residues 484–651). While NMR analysis showed that ABD3 is largely structured in the absence of AD binding (not shown), this domain has neither the solubility nor stability properties required for a full structure determination by NMR. In contrast, ABD2 was tractable for structure determination in its free state. Backbone and side-chain resonance assignments for ^13^C,^15^N-ABD2 were determined with overall completion of 72% (**Table S3**). The chemical shifts predict the secondary structure of ABD2 as consisting of three a-helices (**Fig. 3 and S4B**). A solution structure was calculated using chemical shifts, NOEs, and RDCs as experimental constraints (Methods). The resulting structure is well defined from the NMR constraints, with a backbone rmsd of 0.7 Å for the ordered regions (aa 293–322 and 328–354) in the top 20 of 200 total generated structures (**Fig. 3A**, PDB 6ALY).

**Figure 3.**
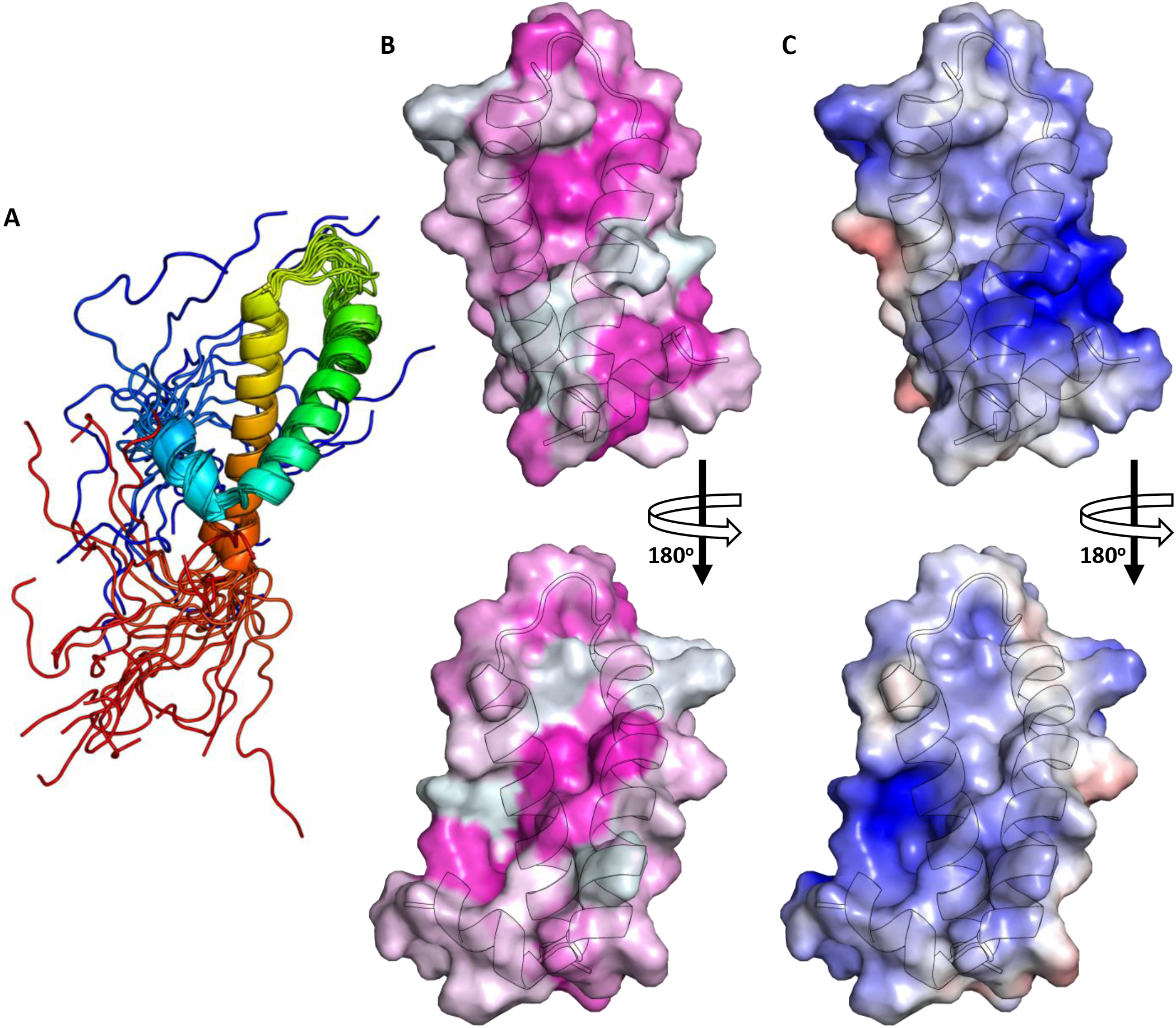
Structure and surface properties of Med15 ABD2. **(A)** The 20 lowest energy structures colored blue to red from the N-to-C-termini. **(B)** Hydrophobic (magenta) and **(C)** electrostatic (red, < −5.0 kT/e; blue, > 5.0 kT/e) surfaces plotted for the front-side and backside of ABD2. Disordered N- and C-termini are not shown for surface representations. Electrostatic surfaces were generated using PDB2QPR and the APBS tool in pymol (Dolinsky et al., 2004). Hydrophobic surfaces are colored from magenta (most hydrophobic) to white according to the Eisenberg hydrophobicity scale.

The tertiary structure of ABD2 is composed of three α-helices; α1 (aa 292–299), α2 (aa 303–323), and α3 (aa 330–354) with the N- and C-terminal tails disordered (**Fig. 3A and S4B**). Unlike ABD1 that binds the cAD via a shallow hydrophobic groove (Brzovic et al., 2011), ABD2 has no obvious groove but instead has two hydrophobic surface patches (**Fig. 3B**). One hydrophobic patch is created at the interface of α2 and α3 and is defined by the near stacking of three aromatic residues (F319, F330, and Y340). A second elongated patch includes α1 and spirals around the ABD2 structure. The ABD2 construct contains 18 charged residues, with a net positive charge of +4. A patch of positive charge separates the two hydrophobic patches (shown as blue, **Fig. 3C**). The structure reveals that although both ABD1 and ABD2 are primarily composed of helices, helical ABDs do not conform to a single structural motif.

### ABD1 and ABD2 use similar strategies to bind ADs of different sequence

To identify the AD-binding site(s) on the ABDs, we performed NMR titration experiments for each combination of ^13^C,^15^N-labeled ABD1 and ABD2 with unlabeled nAD, cAD, and tAD. We showed previously that the cAD binds to ABD1 in a shallow hydrophobic groove as a fuzzy cAD-ABD1 complex (Brzovic et al., 2011). Remarkably, the CSPs in ABD1 are nearly identical regardless of the AD binding partner (**Fig. 4A,C and Fig. S5A**). The similar CSP patterns induced upon binding to each of the three AD variants indicate that their interaction interface is essentially the same. Consistent with this, in a competition binding experiment in which unlabeled cAD was added to ^15^N-tAD + ABD1, all ^15^N-tAD peaks returned towards their chemical shifts in the unbound tAD spectrum, indicating that cAD can compete off both cAD and nAD residues (**Fig. S3C**). The simplest explanation for these combined observations is that ABD1 does not bind the different ADs with a sequence-specific interface, but rather as a “cloud” of hydrophobic character. This model is completely consistent with our earlier characterization of the fuzzy cAD-ABD1 complex.

**Figure 4.**
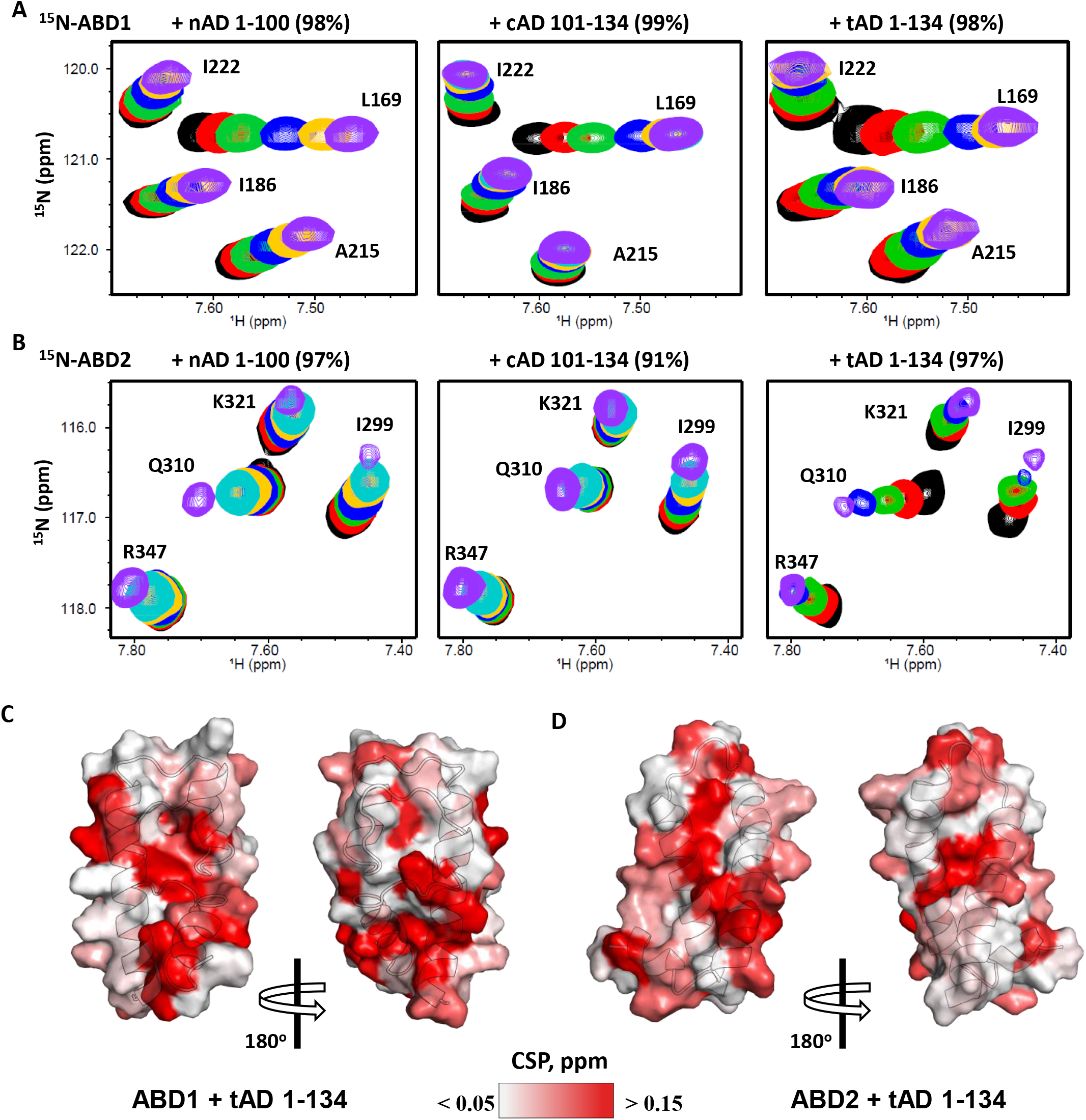
Med15 ABDs show widespread and similar CSPs with each Gcn4 AD. **(A,B)** Expanded region of the (^1^H,^15^N)-HSQC titration spectra of ABD1 **(A)** and ABD2 **(B)** with Gcn4 nAD, cAD, and tAD. Both ABD1 and ABD2 show very similar chemical shift trajectories (examples are shown by black arrows) whether nAD (left), cAD (middle), or tAD (right) is added. Values shown are percent saturation at the end point (purple spectrum). **(C,D)** The CSPs of ABD1 + tAD **(C)** and ABD2 + tAD **(D)** plotted on the structures of ABD1 (2LPB) and ABD2 (6ALY), respectively. A redder surface color indicates a larger CSP.

ABD2 has substantially weaker affinity for cAD than for either nAD or tAD, suggesting that there may be different modes of ABD2 binding. Nevertheless, we again observe highly similar CSPs within ABD2 upon binding the nAD, cAD and tAD peptides (**Fig. 4B,D** and **Fig. S5B**). ABD2 shows widespread CSPs, with the largest perturbations occurring in/near the two ABD2 hydrophobic patches regardless of whether nAD, cAD, or tAD are binding. Therefore, ABD1 and ABD2 each accommodate their AD binding partner(s) in a manner that is independent of the specific sequence of the AD, consistent with a fuzzy binding interface. However, ABD1 contains a single predominant binding groove, while two hydrophobic binding surfaces on ABD2 yield a less localized interaction that may explain the lower affinity for AD-ABD2 binding.

### ABD1 and ABD2 bind ADs with a fuzzy interface

In our previous studies, the definitive evidence for fuzzy binding of cAD-ABD1 derived from NMR spin-label experiments through measurement of paramagnetic relaxation enhancement (PRE) (Brzovic et al., 2011; Warfield et al., 2014). These studies showed that the cAD peptide bound ABD1 in multiple orientations. We performed analogous experiments using a series of nAD, cAD, and tAD constructs to which a single paramagnetic spin-label probe (4-(2-Iodoacetamido)-TEMPO) was attached at sites near, but not in, a primary interacting region. Gcn4 residues T82, T105, and S117 were individually mutated to Cys to allow chemical attachment of the spin label probe (**Fig. 5A**). Groups that are close to a paramagnetic probe will suffer loss of peak intensity, quantified as I_para_/I_dia_ where I_dia_ (diamagnetic) is the reference intensity after reduction of the paramagnetic probe with ascorbic acid. If there is a preferred orientation of an AD in the AD-ABD complex, we expect different patterns of intensity loss when the spin-label is on opposite ends of the AD peptide. We collected (^1^H,^15^N)- and (^1^H,^13^C)-HSQC spectra for ABD1 and ABD2 with each of the spin-labeled ADs at ∼50% saturation and plotted the location of intensity loss for NH and CH groups on the structures of ABD1 and ABD2 (**Fig. 5C** and **5D**, respectively). In every case, the observed patterns of intensity loss are highly similar regardless of which TEMPO-AD was added (**Fig. 5B and S6**). These results indicate that the hydrophobic clusters of the ADs bind without any discernable orientation bias, either when presented as individual ADs, or when both nAD and cAD are present in the tandem construct. Altogether, the results fit a model in which fuzzy binding interfaces are used by each ABD-AD interaction. Since our results above showed that the interactions of the tAD with ABD1 and ABD2 is equivalent in the individual and larger complex, we conclude that the AD-ABD interactions in the larger complex are also fuzzy. Using this mechanism for binding allows the multiple hydrophobic clusters in the tAD to increase binding affinity through avidity, but neither limits nor alters the possible ensemble of complexes that simultaneously exist.

**Figure 5.**
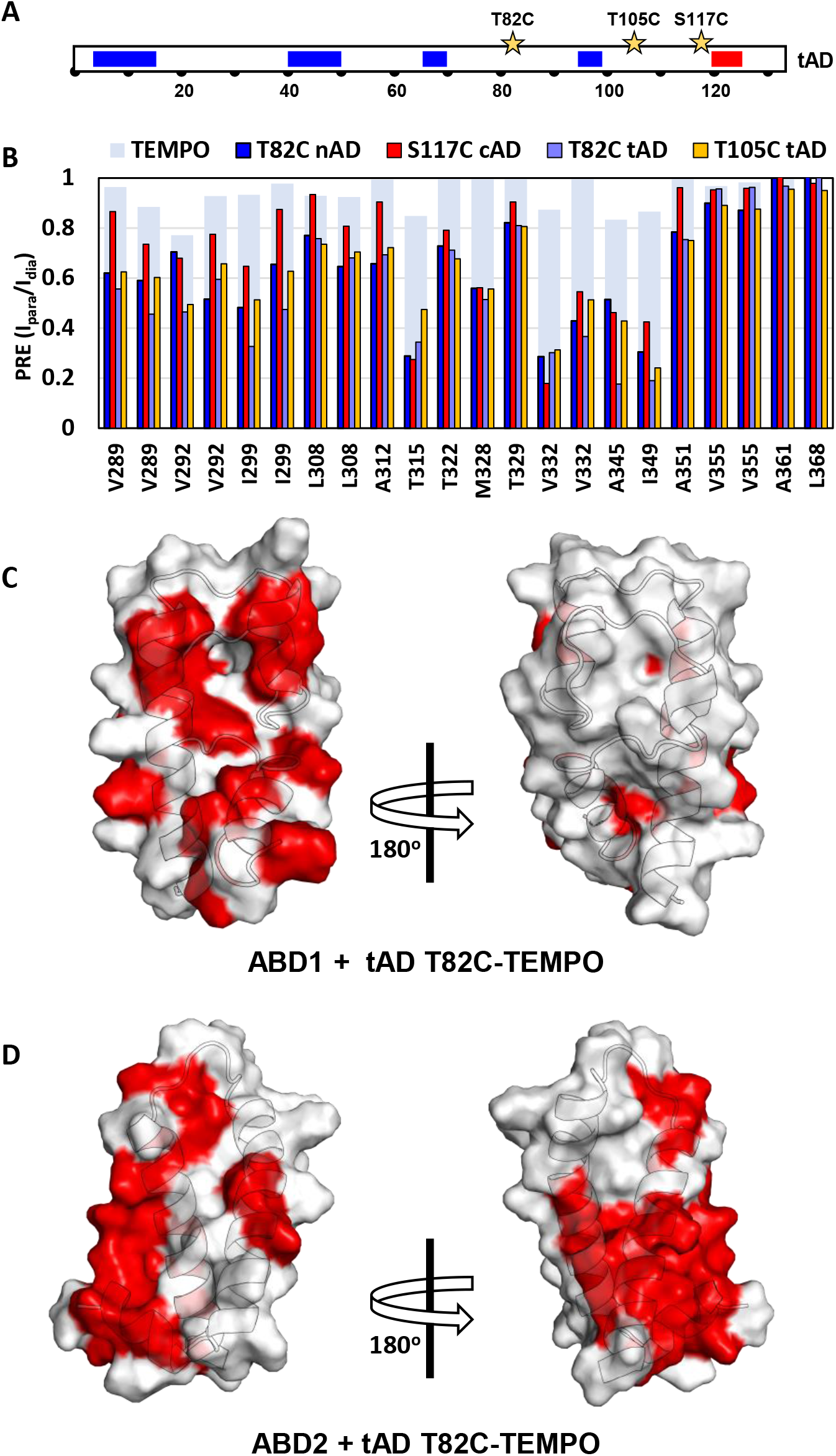
Paramagnetic Relaxation Enhancement experiments of AD:ABD complexes. **(A)** Spin-label was attached at either side of the main clusters of nAD, cAD, and tAD (position indicated by a star). **(B)** Each spin-labeled AD causes similar peak intensity loss at ABD2 (shown here for CH groups) and at ABD1 (see **Fig. S6** for NH and CH results). The results of a control experiment of ABD2 + free TEMPO are shown as light blue bars (see also **Fig. S6F**). **(C,D)** Surface plots for ABD1 + tAD T82C-TEMPO **(C)** and ABD2 + tAD T82C-TEMPO (D). Residues are colored red if the amide PRE < 0.7 or CH group PRE < 0.6.

## Discussion

A longstanding problem in gene regulation has been understanding how many seemingly unrelated transcription activators converge on a limited number of coactivator targets. This situation also raises the question of specificity in activator-coactivator interactions. For example, the fuzzy binding interface of the Gcn4 cAD with Med15 ABD1 has seemingly low specificity and affinity and is not sufficient to efficiently activate transcription by itself (Brzovic et al., 2011; Herbig et al., 2010). Additionally, transcription of many genes requires multiple activators and it is not clear how or why specific combinations of activators synergize. To investigate this long-standing question, we examined the binding of Gcn4 with the Mediator subunit Med15. Gcn4 has the common arrangement of tandem ADs while Med15 has multiple ABDs, typical of most coactivator complexes. Based on earlier findings, two models can explain the requirement for multiple ADs and ABDs in this system: (1) expansion of the cAD-ABD1 fuzzy binding mechanism to encompass dynamic fuzzy interfaces between the two ADs and all ABDs, or (2) a mechanism where the final complex adopts a discrete state due to stable binding and specific AD-ABD interactions. The latter model has been observed in two systems involving IDP binding to structured proteins that contain multiple binding sites (Dyson and Wright, 2016; Saio et al., 2014), while the former model has yet to be observed experimentally. The distinctions between these models has important implications for the flexibility of AD-ABD combinations used in activation of specific genes and in regulatory circuits where activators can combine to give complex patterns of gene regulation. For example, the “locked down” mechanism requires pairing ADs and ABDs in specific combinations as only particular pairings are expected to work together. At the other extreme, activators like Gcn4, Gal4, VP16, and p53 interact with many unrelated coactivators. Use of one or the other mechanism demands a tradeoff between specificity of interactions versus flexibility in regulation of a wide variety of genes with different coactivator requirements.

Our combined results demonstrate unambiguously that Gcn4-Med15 binding uses a fuzzy “free-for all” mechanism where each AD interacts with all ABDs. Protein crosslinking assays show that Gcn4-Med15 is a heterogeneous complex and binding measurements showed that each Gcn4 AD can bind individual ABD1, 2, and 3 domains with micromolar affinity. Importantly, the crosslinking showed that multiple AD-ABD and AD-KIX binding interactions occur in the complex of full-length Gcn4 and Med15. Binding to KIX in the large complex is especially remarkable as Gcn4-KIX binding is undetectable when separated from the rest of Med15. Our NMR results show that both ADs of Gcn4 bind Med15 independently within the large Gcn4-Med15 complex, again consistent with the fuzzy binding mechanism. Importantly, ABD1 and ABD2 have highly similar NMR CSPs regardless of which AD variants is binding. This behavior is not consistent with a specific binding interface and is again in agreement with the fuzzy binding model where the ABDs seem to recognize a “cloud” of hydrophobicity rather than a specific sequence. Finally, multiple spin-label probes positioned on the tAD showed unequivocally that the tAD binds in multiple orientations on both ABD1 and ABD2.

In recent cryo-EM structures of yeast Mediator with and without other PIC components, the Med15-containing tail module remains disordered even when Gcn4 is present (Nozawa et al., 2017; Robinson et al., 2015; Tsai et al., 2017). This is consistent with our findings that Med15 contains multiple well-ordered activator binding domains separated by long flexible linkers and that interactions of Gcn4 with each ABD do not result in a well-ordered complex. Given the sequence variability of the ADs studied here and our previous findings that even more strongly binding synthetic ADs bind with a fuzzy interface (Warfield et al., 2014), we expect that fuzzy binding is a general feature of many large AD-Med15 complexes. Furthermore, the Med15 ABDs may be representative of protein domains designed to accommodate fuzzy interactions.

While both ABD1 and ABD2 bind Gcn4 with a fuzzy interface, each ABD uses a different type of surface feature to bind ADs. ABD1 has a shallow hydrophobic groove that can accommodate a variety of hydrophobic sequences in different orientations (Brzovic et al., 2011; Warfield et al., 2014) while ABD2 uses two separate hydrophobic surface patches. On their own, it is hard to imagine that individual AD-ABD interactions are specific enough for a biological response. However, the combination of ADs and multiple ABDs within Mediator gives rise to binding specificity, higher affinity, and function and allows important contributions to be made by very weak protein-protein interactions (e.g., Gcn4-KIX and cAD-ABD2) (**Fig. 6**). This combinatorial mechanism likely explains why most coactivator complexes examined to date such as Mediator, Swi/Snf, and NuA4/SAGA have multiple ABDs. Even though not every activator has multiple ADs, many ADs bind DNA as dimers and a common regulatory strategy is to require combinations of transcription factors to bind to regulatory regions for efficient activation of transcription. We imagine that this mechanism has the same effect as multiple ADs in a single factor. It has been proposed that the fuzzy binding mechanism leads to higher affinity binding, in part by decreasing the dissociation rate due to multiple independent binding sites on both binding partners (Olsen et al., 2017).

**Figure 6.**
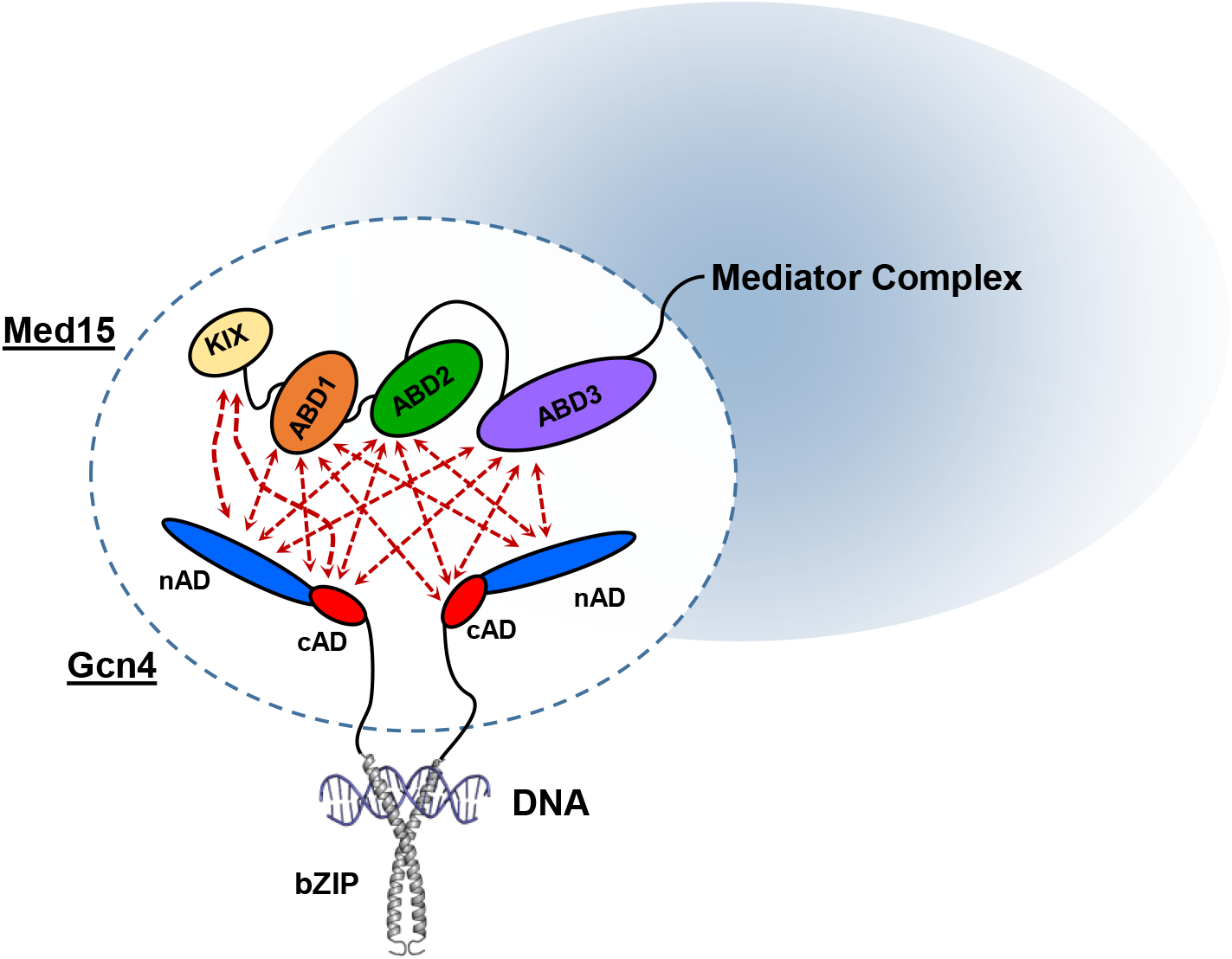
Model of the fuzzy Gcn4-Med15 complex. Transient fuzzy interactions of nAD and cAD with Med15 ABD subunits allows for significant contributions from even the weakest interacting parts by reducing the probability of total dissociation of Gcn4 tAD from Med15. Gcn4 bZIP and DNA structure from PDB: 1DGC.

In the broader context, it is important to note that not all activators work by this mechanism. There are several instances of ADs that bind their targets using specific protein-protein interfaces with much higher affinity. For example, the mammalian coactivator CBP contains numerous activator-binding domains. These individual ABDs partner with specific transcription activators using high affinity conventional protein-protein interfaces (Dames et al., 2002; Demarest et al., 2002; Freedman et al., 2002; Radhakrishnan et al., 1997; Waters et al., 2006; Zor et al., 2004). However, we believe that the fuzzy binding mode is another common regulatory strategy and can account for the behavior in instances where ADs are known to interact with multiple unrelated coactivators. This strategy can generate great flexibility in developing new regulatory circuits where many combinations of activators can regulate a wide variety of genes with different coactivator requirements. It seems likely that a similar strategy will be utilized in other biological systems that utilize IDPs reacting with multiple partners.

## ACKNOWLEDGEMENTS

We thank Ponni Rajagopal and Peter Brzovic of the Klevit laboratory for collection of some of the NMR data. Alexia Loste (Univ Paris, Diderot) for Med15 and Gcn4 expression constructs and for initial crosslinking studies of Gcn4-Med15, and all members of the Klevit and Hahn laboratories for their suggestions and comments during this work. This work was facilitated though the use of advanced computational infrastructure provided by the Hyak supercomputer system at the University of Washington. NMR experiments were performed, in part, in the Environmental Molecular Sciences Laboratories at the Pacific Northwest National Laboratories. This work was funded by NIH RO1 GM075114 to SH and REK and RO1 GM110064 and P50 GM076547 to JR.

## AUTHOR CONTRIBUTIONS

L.M.T., S.H., and R.E.K designed the research. L.M.T, D.P., L.W., J.L., and S.H, performed the experiments. L.M.T., D.P., L.W., J.L., J.R., S.H, and R.E.K analyzed data. L.M.T., D.P., L.W., J.R., S.H, and R.E.K. wrote the paper.

## STAR METHODS

### CONTACT FOR REAGENT AND RESOURCE SHARING

Further information and requests for resources and reagents should be directed to and will be fulfilled by the Lead Contacts, Steve Hahn or Rachel Klevit.

#### Yeast strains and plasmids

Yeast strain SHY823 (*Δgcn4, leu2*) was transformed with the following *LEU2* and GCN4-derivative plasmids for protein expression and mRNA analysis: pRS315 (vector – no *GCN4*), pSH940 (WT *GCN4*), pSH943 (Gcn4 Δ101–124; (nAD) with the flexible linker: GSGSGS at the junction of the internal deletion) (Herbig et al., 2010). Gcn4 nAD mutations were generated in pSH943 by site directed mutagenesis. All Gcn4 derivatives contained a C-terminal 3X-Flag tag.

#### mRNA analysis

RNA was extracted, assayed in triplicate by RT-qPCR, and the results analyzed as described (Herbig et al., 2010).

#### Protein purification

All proteins were expressed in BL21 (DE3) RIL E. coli. Med15 484–651 (ABD3), Med15 158–651 Δ239–272, Δ373–483 (ABD123) and Med15 1–651 Δ239–272, Δ373–483 (KIX123) were expressed as N-terminal His6-tagged proteins. All other Med15 and Gcn4 constructs were expressed as N-terminal His6-SUMO-tagged proteins. Cells were lysed in 50 mM HEPES pH 7.0, 500 mM NaCl, 40 mM Imidazole, 10% glycerol, 1 mM PMSF, 5 mM DTT and purified using Ni-Sepharose High Performance resin (GE Healthcare). Proteins were eluted in 50 mM HEPES pH 7.0, 500 mM NaCl, 500 mM Imidazole, 10% glycerol, 1 mM PMSF, 1 mM DTT. Purified SUMO-tagged proteins were concentrated using 10K MW cutoff centrifugal filters (Millipore), diluted 10x in 50 mM HEPES pH 7.0, 500 mM NaCl, 40 mM Imidazole, 10% glycerol, 1 mM PMSF, 5 mM DTT, and digested with SUMO protease for 3–5 hrs at room temperature using ~1:800 protease:protein ratio. Cleaved His6-Sumo tag was removed using Ni-Sepharose. Med15 polypeptides were further purified using HiTrap Heparin (GE Healthcare) in 20 mM HEPES pH 7, 1 mM DTT, 0.5mM PMSF eluting with either a 50–350 mM NaCl gradient (ABD1 and ABD2) or a 200–600 mM NaCl gradient (ABD3, ABD123, and KIX123). Gcn4 AD derivatives 1–134 and 101–134 were further purified by chromatography on Source 15Q (GE Healthcare) using a 50–350 mM NaCl gradient. Gcn4 1–100 and other nAD derivatives were purified using Source 15Q and HiTrap Phenyl FF (GE Healthcare). Protein was loaded to Source 15Q at 120 mM NaCl and flowed through the column. This unbound fraction was adjusted to 0.8 M (NH4)2SO4, bound to phenyl FF, and eluted with an 800–0 mM (NH4)2SO4 gradient. All proteins were further purified using size exclusion chromatography on Superdex 75 10/30 (GE Healthcare). Proteins used in fluorescence polarization and isothermal titration calorimetry were eluted in 20 mM KH2PO4, pH 7.5, 200 mM KCl. Proteins used in crosslinking-MS were eluted in PBS pH7.2. Proteins used in NMR were eluted in 20 mM NaH2PO4 pH 6.5, 200 mM NaCl, 0.1 mM EDTA, 0.1 mM PMSF, 5 mM DTT. The concentration of the purified proteins was determined by UV/Vis spectroscopy with extinction coefficients calculated with ProtParam (Gasteiger et al., 2005).

#### FP and ITC binding experiments

Gcn4 peptides used in fluorescence polarization were labeled with Oregon Green 488 dye (Invitrogen) as previously described (Herbig et al., 2010). FP measurements were conducted using a Beacon 2000 instrument as previously described (Herbig et al., 2010). Protein concentrations for titrations with Gcn4 nAD and Gcn4 cAD were as described (Herbig et al., 2010), except for Gcn4 cAD vs Med15 ABD3 (0–225 μM). Titrations between Gcn4 tAD and Med15 were performed with 15 concentrations of Med15 spanning 0–135 μM (KIX) or 0–250 μM (ABD1, ABD2, ABD3, ABD123, KIX123). FP data was analyzed using Prism 7 (Graphpad Software, Inc.) to perform non-linear regression analysis using the one-site total binding model Y=B_max_*X/(K_d_+X) + NS*X + Background where Y equals arbitrary polarization units and X equals Med15 concentration.

Isothermal calorimetry titrations were performed using a Microcal ITC200 Microcalorimeter in 20 mM KH_2_PO_4_, pH 7.5, 200 mM KCl as described in (Brzovic et al., 2011). The following protein concentrations were used, with the cell molecule listed first and the syringe molecule listed second: Med15 ABD1 (0.150 mM) vs. Gcn4 nAD (1.50 mM); Gcn4 nAD (0.111 mM) vs. Med15 ABD2 (1.60 mM); Gcn4 nAD (0.111 mM) vs. Med15 ABD3 (1.12 mM); Med15 ABD1 (0.270 mM) vs. Gcn4 cAD (2.37 mM); Med15 ABD2 (0.224 mM) vs. Gcn4 cAD (2.35 mM); Med15 ABD3 (0.236 mM) vs. Gcn4 cAD (2.35 mM). Calorimetric data were plotted and fit with a single binding site model using Origin 7.0 software (Microcal). Some affinities could not be measured by ITC, either due to weak binding or low heats of binding.

#### Protein crosslinking and sample preparation for mass spectrometry

50 μg of Med15 158–651 Δ239–272, Δ373–483 (ABD123) or Med15 1–651 Δ239–272, Δ373–483 (KIX123) was mixed with 3x molar excess of Gcn4 1–140. Samples were incubated with 5 mM (for ABD123 experiments) or 7.5 mM (for KIX123 experiments) EDC (1-ethyl-3-(3-dimethylaminopropyl)carbodiimide hydrochloride; Thermo Scientific), and 2 mM Sulfo-NHS (Thermo Scientific) in 100 μl total volume for 2 hours at room temperature. Protein samples were reduced with 50 mM TCEP and denatured with 8 M urea at 37°C for 15 min. The samples were then alkylated in the dark at 37°C with 15 mM iodoacetamide for 1 hour. The samples were then diluted 10-fold with 100 mM ammonium bicarbonate and digested with trypsin (1:15 w/w) overnight at 37°C. Digested samples were purified by C18 chromatography (Waters), eluted in 80% acetonitrile 0.15 trifluoroacetic acid, and dried in a speedvac.

#### Mass spectrometry for identification of crosslinks

Peptides were analyzed using a Thermo Scientific Orbitrap Elite with HCD fragmentation and serial MS events that included one FTMS1 event at 15,000 resolution followed by 10 FTMS2 events at 7500 resolution. Other instrument settings included: Charge state rejection: +1, +2, +3 and unassigned charges; Monoisotopic precursor selection enabled; Dynamic exclusion enabled: repeat count 1, exclusion list size 500, exclusion duration 30s; HCD normalized collision energy 35%, isolation width 3Da, minimum signal count 5000; MS mass range: > 1500, use m/z values as masses enabled; FTMS MSn AGC target 500,000, FTMS MSn Max ion time 300ms. Peptides were resolved by online reverse phase HPLC using a 90 min gradient from 5% ACN to 40% ACN.

To identify EDC-crosslinked peptides, two different database search algorithms were used: pLink (Yang et al., 2012) and in-house designed Nexus (Luo et al., 2015). pLink was run with default settings (precursor monoisotopic mass tolerance: ± 10 ppm; fragment mass tolerance: ±20 ppm; up to 4 isotopic peaks; max evalue 0.1; static modification on Cysteines; 57.0215 Da; differential oxidation modification on Methionines; 15.9949 Da) using a database containing the target protein sequences. For Nexus searches, a protein database containing the forward and reversed sequences of the target proteins was used with the following parameter settings: up to three miscleavages; static modification on Cysteines (+57.0215 Da); differential oxidation modification on Methionines (+15.9949 Da); differential modification on the peptide N-terminal Glutamic acid residues (-18.0106 Da) or N-terminal Glutamine residues (-17.0265 Da). GluC and trypsin were specified as the digestion enzymes, and a 5% of FDR was used for both searches. After performing the pLink and the Nexus analysis, the search results are combined and each spectrum was manually evaluated for the quality of the match to each peptide using the COMET/Lorikeet Spectrum Viewer (TPP). A crosslinked peptide is considered to be confidently identified if at least four consecutive b or y ions for each peptide is observed and the majority of the observed ions are accounted for.

#### NMR experiments and resonance assignments

NMR HSQC titration and spin-label experiments were completed on a Bruker 500 MHz AVANCE spectrometer. All spectra were collected at 25 °C in NMR buffer (20 mM sodium phosphate, 150 mM NaCl, 0.1 mM EDTA, 0.1 mM PMSF, and 5 mM DTT) with 7% D_2_O, unless otherwise specified. Spin-label samples were in NMR buffer but with no DTT; titration samples were in NMR buffer, but with 200 mM NaCl because ABD3 is more stable at higher salt.

(^1^H,^15^N)- and (^1^H,^13^C)-HSQC titration experiments were completed for all Gcn4-AD:Med15-ABD combinations by adding unlabeled AD (nAD, cAD, tAD) or ABD (ABD1, ABD2, ABD3, ABD123) to a ∼200 μM [^13^C,^15^N]-AD or ABD sample, maintaining a constant concentration of the labeled species. Paramagnetic relaxation enhancement (PRE) experiments were performed as for cAD:ABD1 (Brzovic et al., 2011), but at lower saturation. The spin-label 4-(2-Iodoacetamido)-TEMPO was incorporated at T82C (nAD and tAD), T105C (tAD), and S117C (cAD and tAD) by incubating the AD Cys mutants with 10x TEMPO overnight at room temperature, followed by several hours at 30 °C. Excess TEMPO was removed by elution over a Nap-10 column and buffer exchange during concentration. ^15^N- and ^13^C-HSQC spectra of [^13^C,^15^N]-ABD1 and [^13^C,^15^N]-ABD2 with spin-labeled ADs at a ∼2:1 ABD:AD concentration ratio were collected in the presence (reference intensity, Idia) and absence of 3 mM ascorbic acid (Ipara). A control sample was collected of [^13^C,^15^N]-ABD2 with 500 μM free TEMPO, with and without 3 mM ascorbic acid.

Gcn4 tAD (aa 1–134) chemical shift assignments were transferred from nAD (aa 1–100) and cAD (aa 101–134) assignments (Brzovic et al., 2011) where possible, and verified by standard backbone triple-resonance experiments (HNCA, HNCOCA, HNCOCACB, HNCACB, and HNCO) obtained on a 400 μM [^13^C,^15^N]-tAD sample using Bruker 500 and 600 MHz Avance spectrometers. Gcn4 nAD chemical shift assignments were transferred from mini-nAD (Gcn4 1–100, Δ21–40, 52–60) where possible, and verified by standard backbone triple-resonance experiments (HNCA, HNCOCA, HNCOCACB, HNCACB, HNCO) obtained on a 350 μM [^13^C,^15^N]-nAD + 620 μM ABD1 sample using Bruker 500 and 600 MHz Avance spectrometers. Assignments of free nAD were determined by following HSQC titration trajectories of nAD + ABD1. Mini-nAD assignments were obtained on a 600 μM [^13^C,^15^N]-mini-nAD sample using standard experiments (HNCA, HNCOCA, HNCACB, HNCOCACB, HNCO) on a Varian INOVA 600 or 800 MHz instruments located at Pacific Northwest National Laboratories.

Med15 ABD2 (residues 277–368) backbone and side-chain chemical shift assignments were obtained on ∼1 mM [^13^C,^15^N]-ABD2 using standard experiments (HNCA, HNCOCA, HNCACB, HNCO, ^15^N-TOCSY, ^15^N-NOESY, aromatic and aliphatic ^13^C-NOESY, HNHAHB, HCCONH, CCONH, HCCH-TOCSY, HCCH-COSY) on Bruker 600 and 800 MHz Avance spectrometers. ^13^C-NOESY and HCCH experiments were collected in D2O NMR buffer. HSQC and NOESY experiments were also collected on a [^13^C,^15^N]-F319Y ABD2 mutant sample to assist in assignments of aromatic residues and NOEs (**Fig. S4A**). (^1^H,^15^N)-HSQC-IPAP experiments for measuring D_NH_ Residual Dipolar Couplings (RDCs) were collected on a ^15^N-ABD2 sample at 22 °C using a C12E6/hexanol mixture for alignment (Higman et al., 2011) on a Bruker 800 MHz Avance spectrometer.

The NH chemical shift perturbation was calculated according to Δδ_NH_ (ppm) = sqrt [Δδ_H^2^_ + (Δδ_N_/5)^2^]. Secondary structure propensity was determined from backbone chemical shifts using the Neighbor Corrected Structural Propensity Calculator (ncSPC) (Tamiola and Mulder, 2012). Max CSP as in **Figure S3A** represents the maximum CSP of any residue within each defined hydrophobic cluster.

#### NMR solution structure

The structure of free-form Med15 ABD2 (aa 277–368) was determined based on chemical shifts, NOEs, and RDCs using the Xplor-NIH software (Schwieters et al., 2003) with the EEFx implicit water potential (Tian et al., 2014). NOEs were assigned manually and used as distance restraints. Dihedral backbone restraints were calculated from backbone chemical shifts using TALOS (Shen and Bax, 2013) and were only used for residues with ‘Strong’ database matches. Structure calculations were based on the recommended eefx/fold.py and refine.py scripts, which are included with the software. Simulations start from an extended conformation of ABD2, followed by rounds of simulated annealing and cooling to find the 20 lowest energy of 200 generated structures. Structure validation and statistics (**Table S3**) were determined using the Protein Structure Validation Software (PSVS) web server (Basu et al., 2008) and Procheck via the wwPDB submission (Berman et al., 2003). Electrostatic surfaces were generated using PDB2QPR and the APBS tool in pymol (Dolinsky et al., 2004).

### QUANTIFICATION AND STATISTICAL ANALYSIS

#### mRNA quantification

Steady-state mRNA levels were quantitated by RT qPCR as described above and in (Herbig et al., 2010).

#### Mass spectrometry and data analysis

EDC–cross-linked peptides were analyzed on a Thermo Scientific Orbitrap Elite at the Proteomics facility at the Fred Hutchinson Cancer Research Center and data were analyzed as described(Knutson et al., 2014). Two different database search algorithms were used to identify EDC-crosslinked peptides; pLink (Yang et al., 2012) and the in-house designed Nexus (Luo et al., 2015). A 5% of FDR was used for the results from both searches. Spectra were manually evaluated using the COMET/Lorikeet Spectrum Viewer (Trans-Proteomic Pipeline) (Knutson et al., 2014).

#### NMR analysis

All NMR spectra were processed using NMRPipe (Delaglio et al., 1995), and analyzed in nmrViewJ (Johnson, 2004). NMR peak intensities for spin-label experiments were quantified using NMRviewJ and error bars reflect noise levels of the spectra. Peak centers for RDC measurements were determined using FuDA (Hansen et al., 2007). Percent saturation was calculated based on the K_d_ for each A:B complex according to %sat = [AB]/A = (1/A) *(C/2+(C^2^ – 4*A*B)^0.5^), where A=[A_total_], B=[B_total_], and C=A+B+K_d_.

## DATA AND SOFTWARE AVAILABILITY

The solution structure of Med15 ABD2 has been deposited in the Protein Data Bank with the accession code 6ALY. NMR chemical shifts, NOEs, and RDCs have been deposited to the BioMagResBank: ABD2 (accession number 30330), Gcn4 tAD 1–134 (accession number 27207). NMR and cross-linking data has been deposited in the Mendeley Data repository (http://dx.doi.org/10.17632/twjfm5rnnm.1).

## KEY RESOURCES TABLE

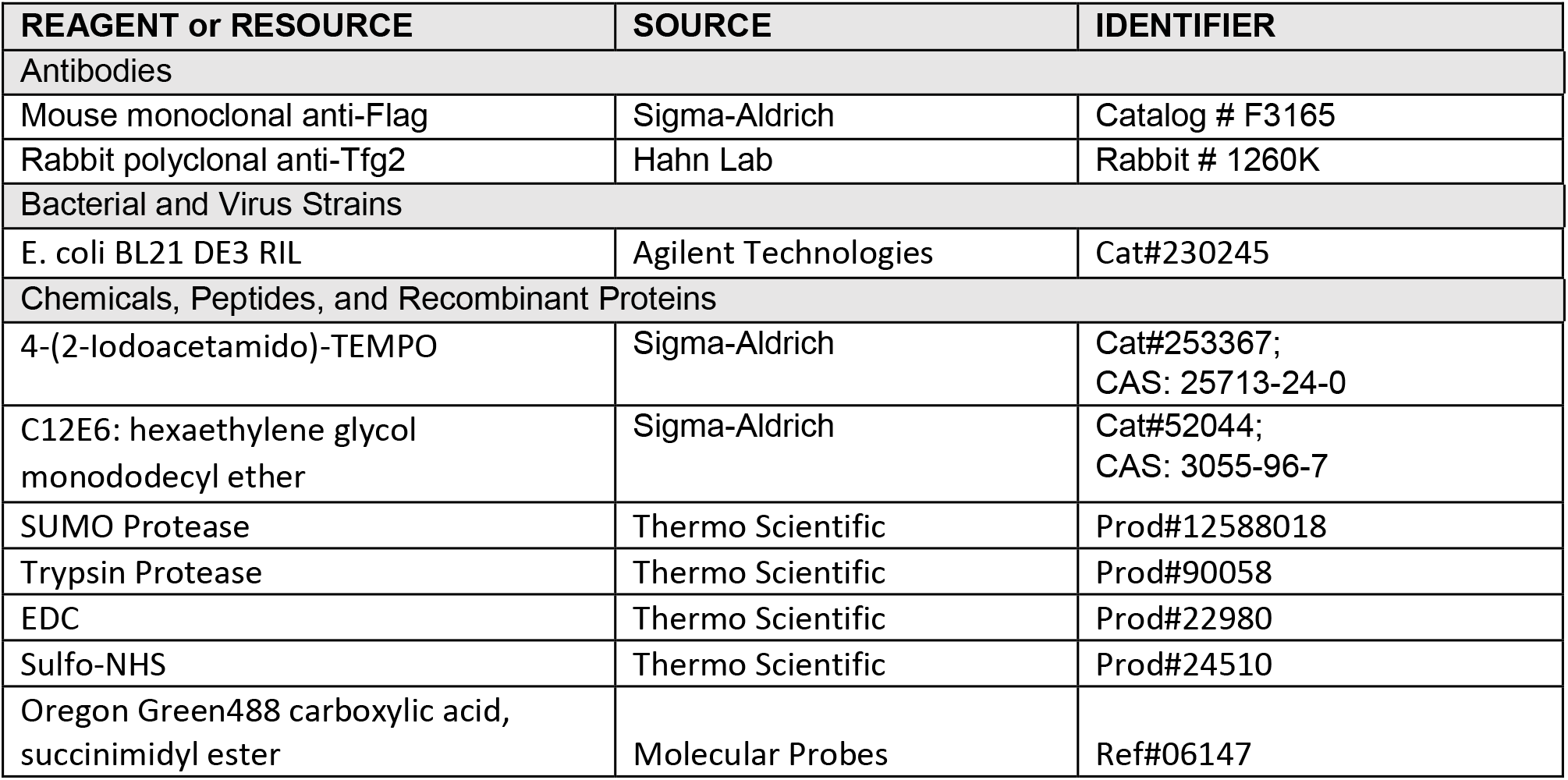
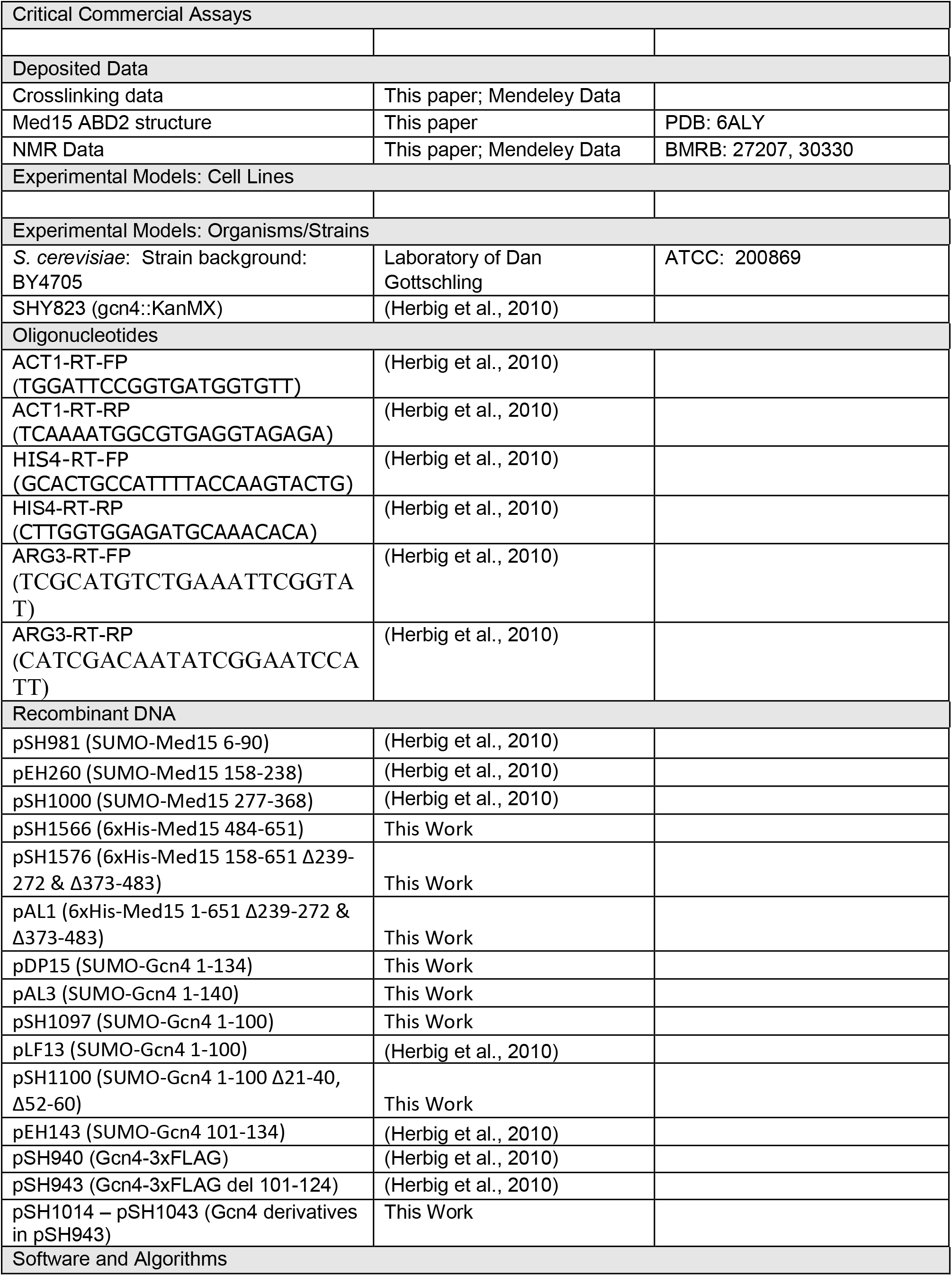
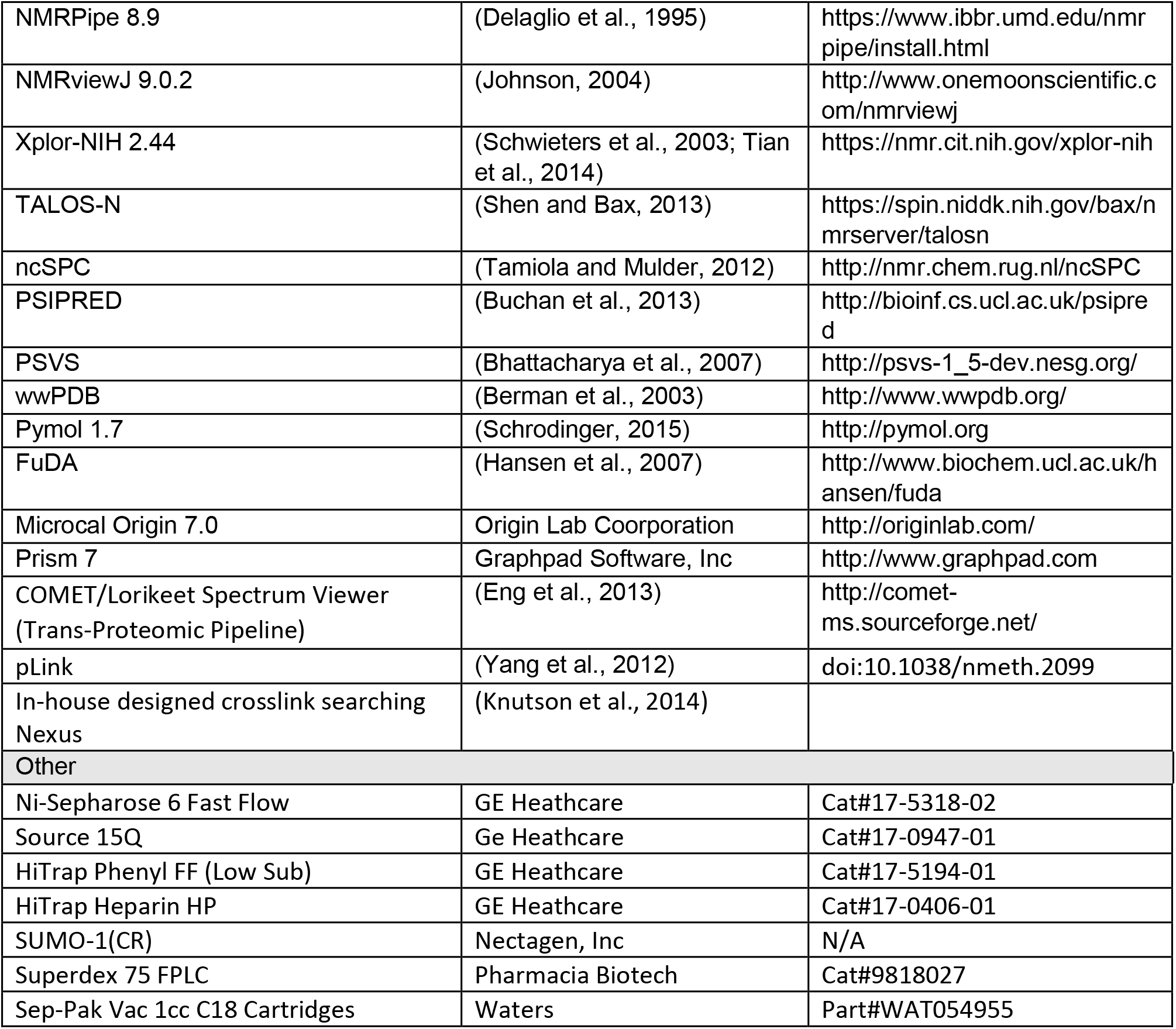

---

**Figure S1.**
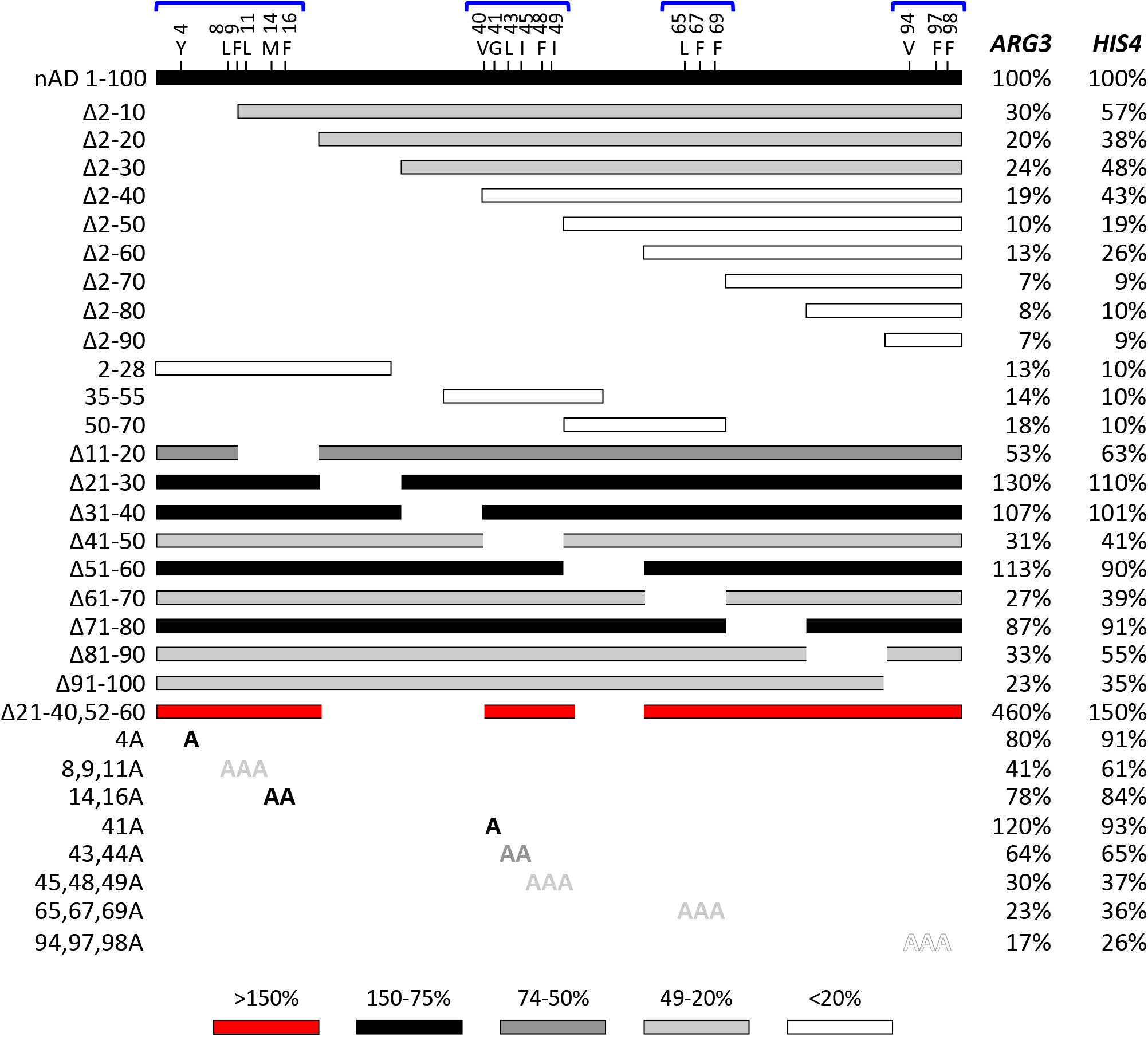
Gcn4 nAD mutations reveal regions critical for transcription activity. Activation of transcription of *ARG3* and *HIS4* in yeast by Gcn4 nAD derivatives. Each isolated hydrophobic region of the Gcn4 nAD is only weakly activating and deletion of any of these regions decreases *ARG3* transcription activation levels ≥ 3-fold. Shading of bars and alanine substitutions indicates activation function relative to wild-type nAD.

**Figure S2.**
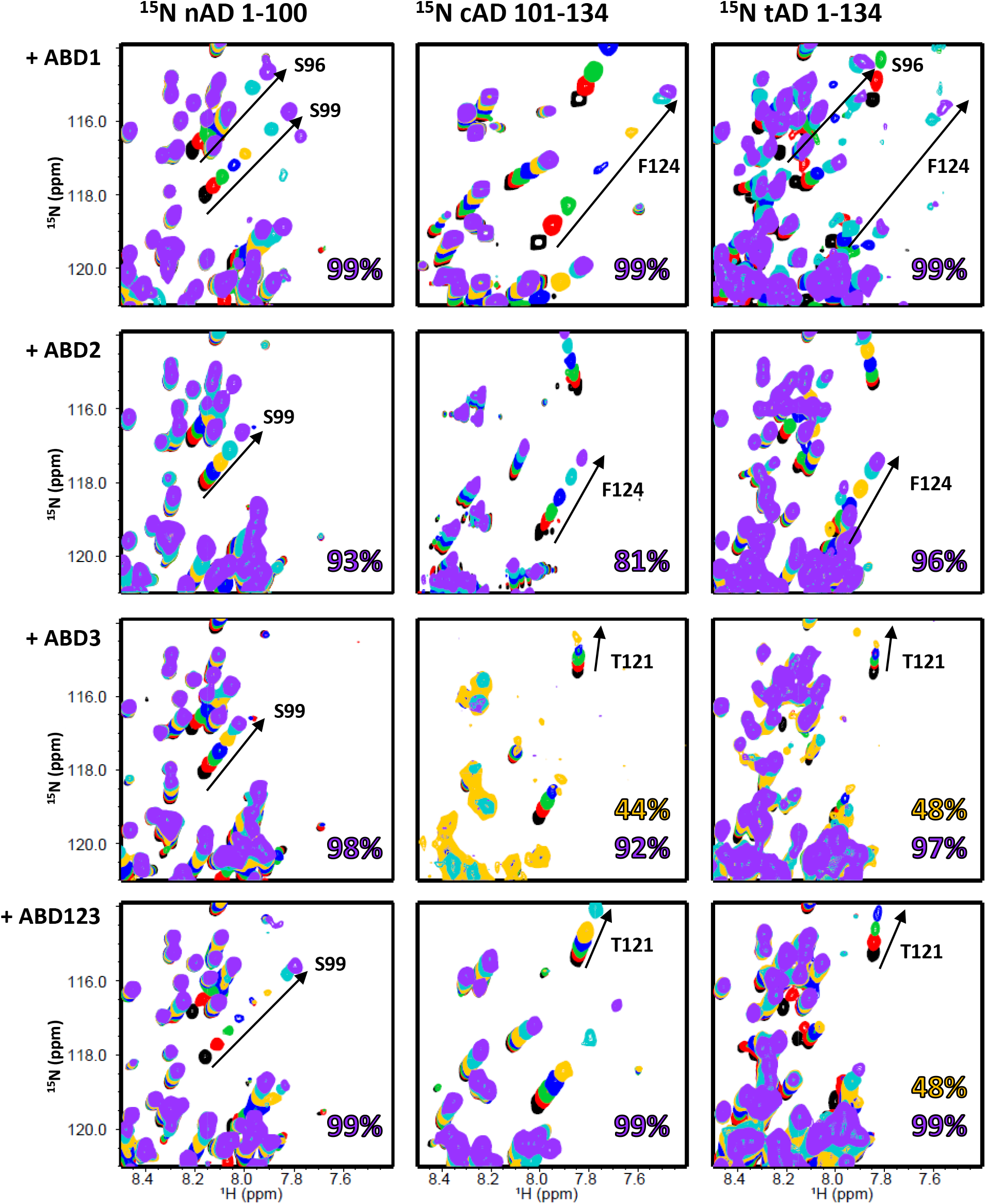
Related to Figure 2. Region of the (^1^H,^15^N)-HSQC overlaid titration spectra of labeled nAD, cAD, and tAD with unlabeled ABD1, ABD2, ABD3, and ABD123. Titrations are from free-form AD (black) to saturated AD:ABD (purple). Numbers on each plot represent the percent saturation of the last visible titration point based on Kd values given in Table 1. Some peaks in tAD+ABD123, cAD+ABD3, and tAD+ABD3 are lost after ~50% saturation. Contours are chosen for clarity. Residues 98 and 99 of nAD 1–100 have slightly different chemical shifts in the context of tAD 1–134, but show nearly identical shift trajectories in nAD and tAD upon addition of ABD. Positions of cAD shifting peaks are unchanged in tAD 1–134.

**Figure S3.**
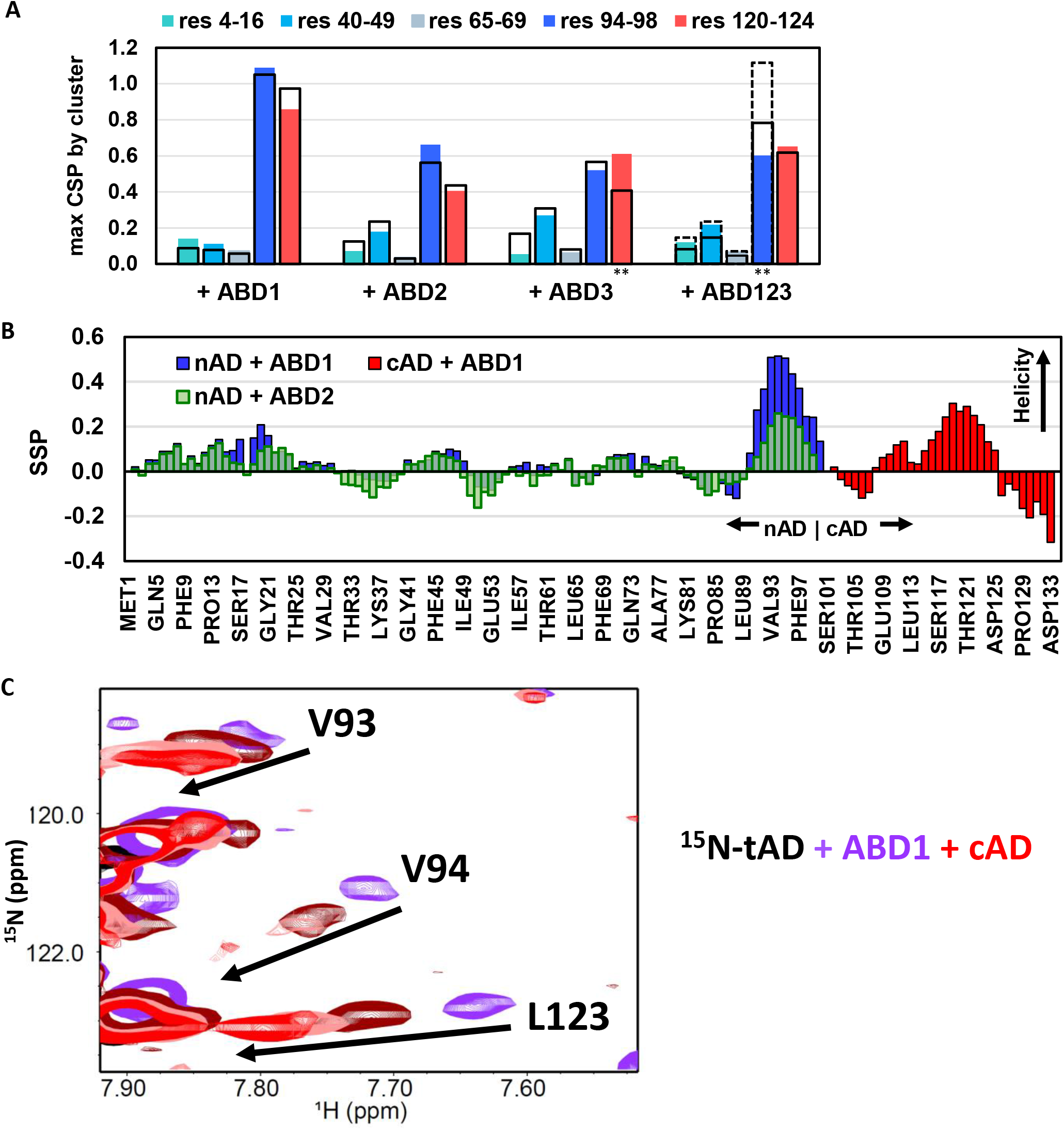
Related to Figure 2. Characteristics of AD + ABD binding. **(A)** Contribution of each AD hydrophobic cluster to ABD binding. Filled bars represent the maximum CSP value of any residue in each cluster for tAD + ABD titrations; black borders are used for nAD + ABD and cAD + ABD titrations. Max CSP for cluster 94–98 of nAD +ABD123 and tAD + ABD123 represents a ~50% saturation, dashed black borders are for full saturation nAD + ABD123. ** indicate situations where not all peak trajectories could be followed to complete saturation. **(B)** Secondary Structure Propensity (SSP) for nAD upon binding to ABD1 (blue) or ABD2 (green outlined bars) and for cAD upon binding to ABD1 (red), determined from backbone chemical shifts using the ncSPC webserver. Positive values indicate helical structure; negative values indicate extended structure. **(C)** Expanded region of the (^1^H,^15^N)-HSQC of ^15^N-tAD + ABD1 + cAD showing that cAD can compete off all clusters of tAD. This supports a single binding surface for AD on ABD1.

**Figure S4.**
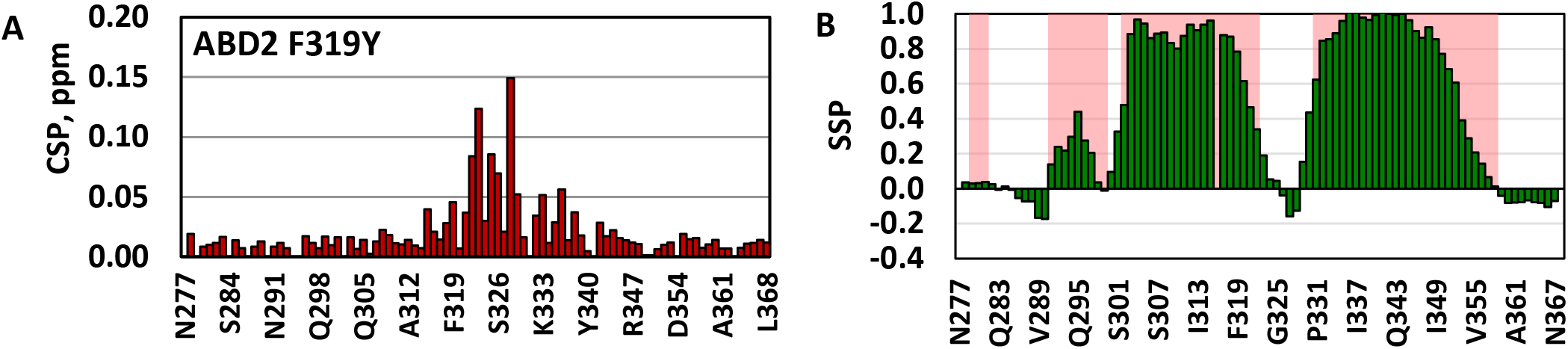
Related to Figure 3. Structure and surface properties of Med15 ABD2. **(A)** The ABD2 F319Y mutation, which was used to assist with aromatic NOE assignments, minimally perturbs the ABD2 ^15^N-HSQC spectra. (B) Green bars show secondary structure propensity for Med15 ABD2 based on backbone chemical shifts calculated using ncSPC. Pink shading indicates regions predicted to be helical based on the amino acid sequence using PSIPRED (Buchan et al., 2013).

**Figure S5.**
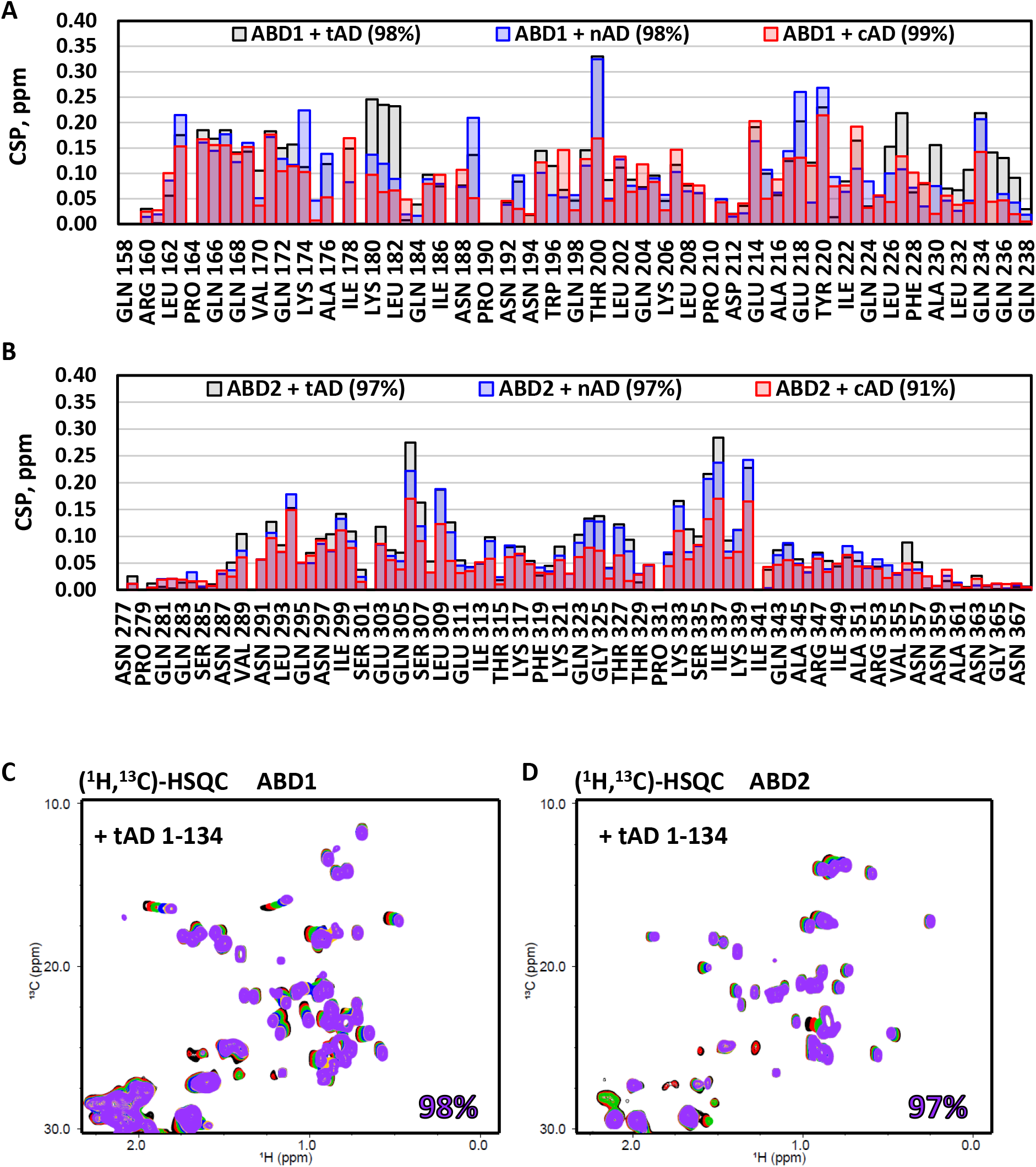
Related to Figure 4. ABD + AD titrations. **(A)** ABD1 and **(B)** ABD2 show widespread NH chemical shift perturbations (CSPs) with each AD. Even cAD, which only weakly binds to ABD2, shows similar CSPs as nAD and tAD. Percent saturation for each titration is given in parentheses in the legend entry. **(C,D)** Methyl region of the (^1^H,^13^C)-HSQC for ABD1 and ABD2 + tAD 1 -134 titrations. Values shown are the percent saturation of the ABD:AD complex at the final titration point, based on the K_d_ values in Table 1.

**Figure S6.**
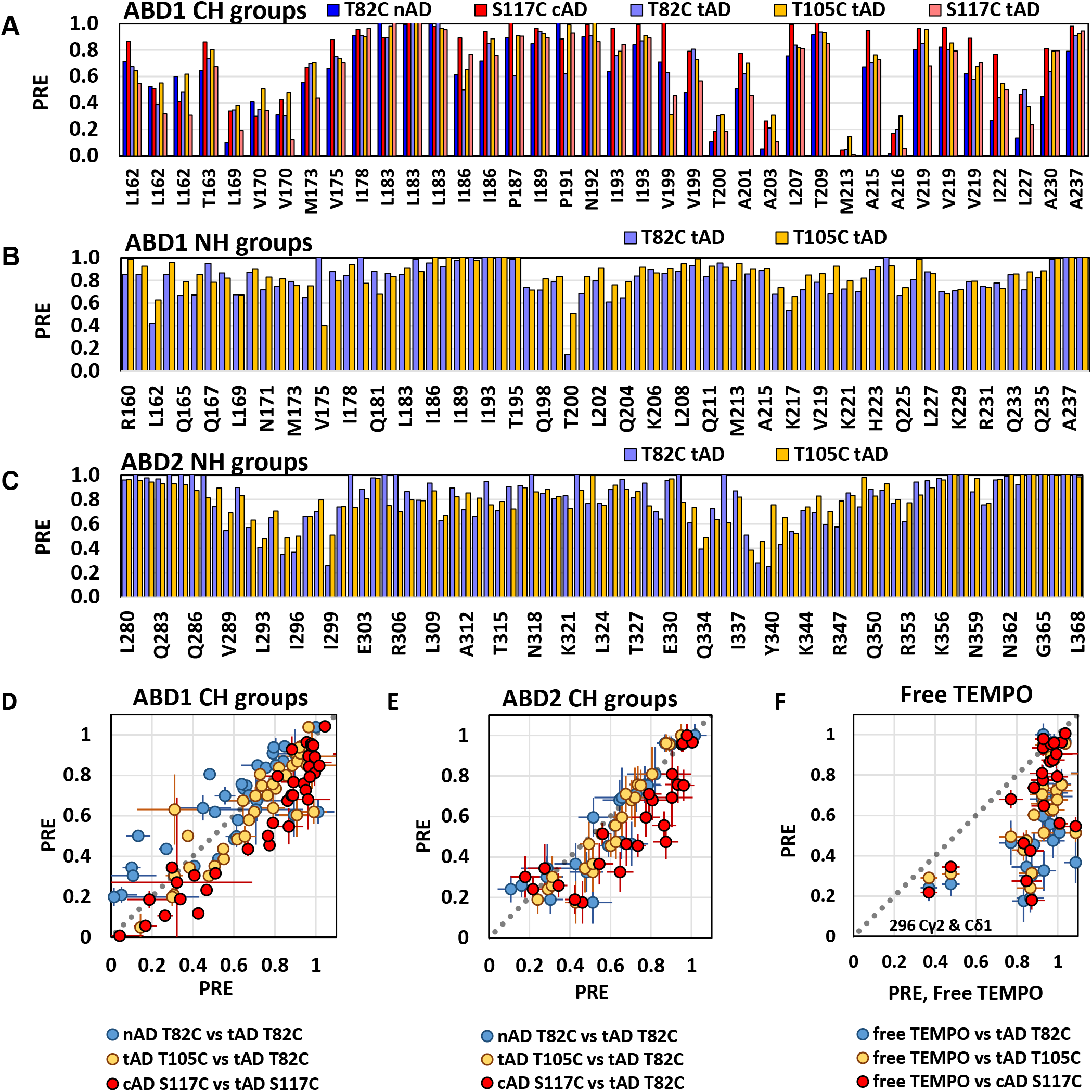
Related to Figure 5. Gcn4 nAD, cAD, and tAD spin labels similarly affect ABD1 and ABD2. **(A)** Paramagnetic Relaxation Effect (PRE) for CH groups of each spin-labeled AD on ABD1. **(B,C)** PRE for NH groups for select spin-labeled ADs for ABD1 **(B)** and ABD2 **(C)**. The magnitude of the PRE depends on saturation levels of each AD:ABD complex. **(D-F)** Scatter plots represent pairwise comparison of the PRE intensity ratio of various AD-TEMPO spin-label experiments. For both ABD1 **(D)** and ABD2 **(E)**, intensity losses are highly correlated regardless of where in nAD, cAD, or tAD the TEMPO spin label is located (magnitude of PRE is dependent on saturation). **(F)** Scatter plot of intensity loss for ABD2 + spin-labeled ADs compared to a control experiment of ABD2 + free TEMPO. Most ABD2 residues show minimal intensity loss upon addition of free TEMPO, supporting that intensity loss in the spin-label AD experiments is due to the AD and not a non-specific effect of the TEMPO probe. Only lle296 methyl groups show significant intensity loss from free TEMPO. A dashed y=x line in **E-G** is shown for reference. Error bars are based on noise levels of each spectra.

**Table S3.**
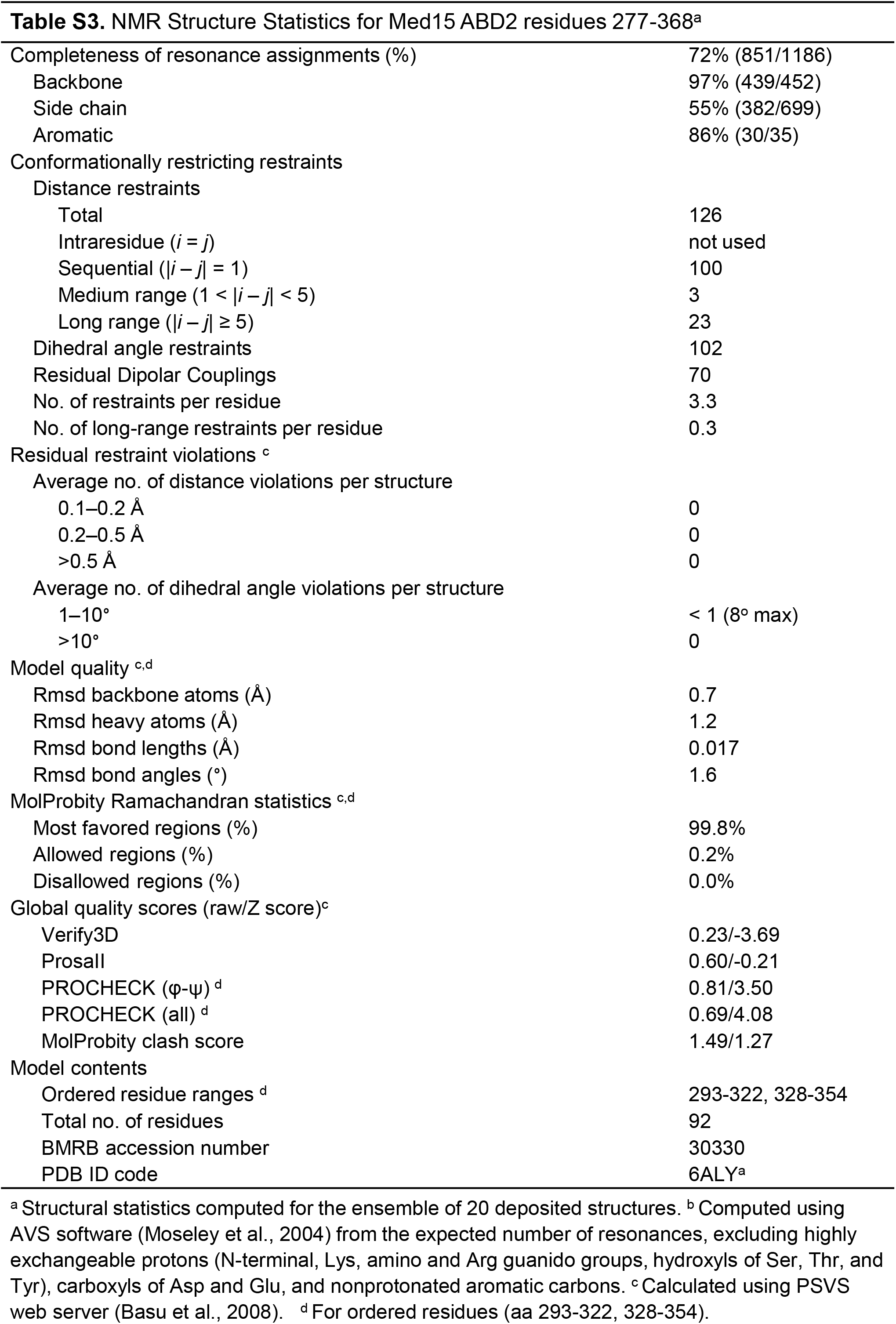
NMR Structure Statistics for Med15 ABD2 residues 277–368^a^

## REFERENCES

Basu, A., Samanta, D., Bhattacharya, A., Das, A., Das, D., and Dasgupta, C. (2008). Protein folding following synthesis in vitro and in vivo: association of newly synthesized protein with 50S subunit of E. coli ribosome. Biochem Biophys Res Commun 366, 592–597.

Berman, H., Henrick, K., and Nakamura, H. (2003). Announcing the worldwide Protein Data Bank. Nat Struct Biol 10, 980.

Bhattacharya, A., Tejero, R., and Montelione, G.T. (2007). Evaluating protein structures determined by structural genomics consortia. Proteins 66, 778–795.

Brown, C.E., Howe, L., Sousa, K., Alley, S.C., Carrozza, M.J., Tan, S., and Workman, J.L. (2001a). Recruitment of HAT complexes by direct activator interactions with the ATM-related Tra1 subunit. Science 292, 2333–2337.

Brown, D.R., Deb, D., Frum, R., Hickes, L., Munoz, R., Deb, S., and Deb, S.P. (2001b). The human oncoprotein MDM2 uses distinct strategies to inhibit transcriptional activation mediated by the wild-type p53 and its tumor-derived mutants. International journal of oncology 18, 449–459.

Brzovic, P.S., Heikaus, C.C., Kisselev, L., Vernon, R., Herbig, E., Pacheco, D., Warfield, L., Littlefield, P., Baker, D., Klevit, R.E., et al. (2011). The acidic transcription activator Gcn4 binds the mediator subunit Gal11/Med15 using a simple protein interface forming a fuzzy complex. Mol Cell 44, 942–953.

Buchan, D.W., Minneci, F., Nugent, T.C., Bryson, K., and Jones, D.T. (2013). Scalable web services for the PSIPRED Protein Analysis Workbench. Nucleic Acids Res 41, W349–357.

Chang, J., Kim, D.H., Lee, S.W., Choi, K.Y., and Sung, Y.C. (1995). Transactivation ability of p53 transcriptional activation domain is directly related to the binding affinity to TATA-binding protein. J Biol Chem 270, 25014–25019.

Dames, S.A., Martinez-Yamout, M., De Guzman, R.N., Dyson, H.J., and Wright, P.E. (2002). Structural basis for Hif-1 alpha /CBP recognition in the cellular hypoxic response. Proc Natl Acad Sci U S A 99, 5271–5276.

Delaglio, F., Grzesiek, S., Vuister, G.W., Zhu, G., Pfeifer, J., and Bax, A. (1995). NMRPipe: a multidimensional spectral processing system based on UNIX pipes. J Biomol NMR 6, 277–293.

Demarest, S.J., Martinez-Yamout, M., Chung, J., Chen, H., Xu, W., Dyson, H.J., Evans, R.M., and Wright, P.E. (2002). Mutual synergistic folding in recruitment of CBP/p300 by p160 nuclear receptor coactivators. Nature 415, 549–553.

Diaz-Santin, L.M., Lukoyanova, N., Aciyan, E., and Cheung, A.C. (2017). Cryo-EM structure of the SAGA and NuA4 coactivator subunit Tra1 at 3.7 angstrom resolution. Elife 6.

Dolinsky, T.J., Nielsen, J.E., McCammon, J.A., and Baker, N.A. (2004). PDB2PQR: an automated pipeline for the setup of Poisson-Boltzmann electrostatics calculations. Nucleic Acids Res 32, W665–667.

Drysdale, C.M., Duenas, E., Jackson, B.M., Reusser, U., Braus, G.H., and Hinnebusch, A.G. (1995). The transcriptional activator GCN4 contains multiple activation domains that are critically dependent on hydrophobic amino acids. Mol Cell Biol 15, 1220–1233.

Dyson, H.J., and Wright, P.E. (2005). Intrinsically unstructured proteins and their functions. Nat Rev Mol Cell Biol 6, 197–208.

Dyson, H.J., and Wright, P.E. (2016). Role of Intrinsic Protein Disorder in the Function and Interactions of the Transcriptional Coactivators CREB-binding Protein (CBP) and p300. J Biol Chem 291, 6714–6722.

Eng, J.K., Jahan, T.A., and Hoopmann, M.R. (2013). Comet: An open-source MS/MS sequence database search tool. PROTEOMICS 13, 22–24.

Ferreon, J.C., Lee, C.W., Arai, M., Martinez-Yamout, M.A., Dyson, H.J., and Wright, P.E. (2009). Cooperative regulation of p53 by modulation of ternary complex formation with CBP/p300 and HDM2. Proc Natl Acad Sci U S A 106, 6591–6596.

Fishburn, J., Mohibullah, N., and Hahn, S. (2005). Function of a eukaryotic transcription activator during the transcription cycle. Mol Cell 18, 369–378.

Freedman, S.J., Sun, Z.Y., Poy, F., Kung, A.L., Livingston, D.M., Wagner, G., and Eck, M.J. (2002). Structural basis for recruitment of CBP/p300 by hypoxia-inducible factor-1 alpha. Proc Natl Acad Sci U S A 99, 5367–5372.

Gasteiger, E., Hoogland, C., Gattiker, A., Duvaud, S., Wilkins, M.R., Appel, R.D., and Bairoch, A. (2005). Protein Identification and Analysis Tools on the ExPASy Server. In The Proteomics Protocols Handbook, J.M. Walker, ed. (Humana Press), pp. 571–607

Hahn, S., and Young, E.T. (2011). Transcriptional regulation in Saccharomyces cerevisiae: transcription factor regulation and function, mechanisms of initiation, and roles of activators and coactivators. Genetics 189, 705–736.

Hansen, D.F., Yang, D., Feng, H., Zhou, Z., Wiesner, S., Bai, Y., and Kay, L.E. (2007). An exchange-free measure of 15N transverse relaxation: an NMR spectroscopy application to the study of a folding intermediate with pervasive chemical exchange. J Am Chem Soc 129, 11468–11479.

Herbig, E., Warfield, L., Fish, L., Fishburn, J., Knutson, B.A., Moorefield, B., Pacheco, D., and Hahn, S. (2010). Mechanism of Mediator recruitment by tandem Gcn4 activation domains and three Gal11 activator-binding domains. Mol Cell Biol 30, 2376–2390.

Higman, V.A., Boyd, J., Smith, L.J., and Redfield, C. (2011). Residual dipolar couplings: are multiple independent alignments always possible? J Biomol NMR 49, 53–60.

Jackson, B.M., Drysdale, C.M., Natarajan, K., and Hinnebusch, A.G. (1996). Identification of seven hydrophobic clusters in GCN4 making redundant contributions to transcriptional activation. Mol Cell Biol 16, 5557–5571.

Jedidi, I., Zhang, F., Qiu, H., Stahl, S.J., Palmer, I., Kaufman, J.D., Nadaud, P.S., Mukherjee, S., Wingfield, P.T., Jaroniec, C.P., et al. (2010). Activator Gcn4 employs multiple segments of Med15/Gal11, including the KIX domain, to recruit mediator to target genes in vivo. J Biol Chem 285, 2438–2455.

Johnson, B.A. (2004). Using NMRView to visualize and analyze the NMR spectra of macromolecules. Methods Mol Biol 278, 313–352.

Knutson, B.A., Luo, J., Ranish, J., and Hahn, S. (2014). Architecture of the Saccharomyces cerevisiae RNA polymerase I Core Factor complex. Nat Struct Mol Biol 21, 810–816.

Langlois, C., Mas, C., Di Lello, P., Jenkins, L.M., Legault, P., and Omichinski, J.G. (2008). NMR structure of the complex between the Tfb1 subunit of TFIIH and the activation domain of VP16: structural similarities between VP16 and p53. J Am Chem Soc 130, 10596–10604.

Levine, M., Cattoglio, C., and Tjian, R. (2014). Looping back to leap forward: transcription enters a new era. Cell 157, 13–25.

Luo, J., Cimermancic, P., Viswanath, S., Ebmeier, C.C., Kim, B., Dehecq, M., Raman, V., Greenberg, C.H., Pellarin, R., Sali, A., et al. (2015). Architecture of the Human and Yeast General Transcription and DNA Repair Factor TFIIH. Mol Cell 59, 794–806.

Natarajan, K., Meyer, M.R., Jackson, B.M., Slade, D., Roberts, C., Hinnebusch, A.G., and Marton, M.J. (2001). Transcriptional profiling shows that Gcn4p is a master regulator of gene expression during amino acid starvation in yeast. Mol Cell Biol 21, 4347–4368.

Nguyen Ba, A.N., Yeh, B.J., van Dyk, D., Davidson, A.R., Andrews, B.J., Weiss, E.L., and Moses, A.M. (2012). Proteome-wide discovery of evolutionary conserved sequences in disordered regions. Sci Signal 5, rs1.

Novatchkova, M., and Eisenhaber, F. (2004). Linking transcriptional mediators via the GACKIX domain super family. Curr Biol 14, R54–55.

Nozawa, K., Schneider, T.R., and Cramer, P. (2017). Core Mediator structure at 3.4 A extends model of transcription initiation complex. Nature 545, 248–251.

Olsen, J.G., Teilum, K., and Kragelund, B.B. (2017). Behaviour of intrinsically disordered proteins in protein-protein complexes with an emphasis on fuzziness. Cell Mol Life Sci.

Prochasson, P., Florens, L., Swanson, S.K., Washburn, M.P., and Workman, J.L. (2005). The HIR corepressor complex binds to nucleosomes generating a distinct protein/DNA complex resistant to remodeling by SWI/SNF. Genes Dev 19, 2534–2539.

Qiu, H., Chereji, R.V., Hu, C., Cole, H.A., Rawal, Y., Clark, D.J., and Hinnebusch, A.G. (2016). Genome-wide cooperation by HAT Gcn5, remodeler SWI/SNF, and chaperone Ydj1 in promoter nucleosome eviction and transcriptional activation. Genome research 26, 211–225.

Radhakrishnan, I., Perez-Alvarado, G.C., Parker, D., Dyson, H.J., Montminy, M.R., and Wright, P.E. (1997). Solution structure of the KIX domain of CBP bound to the transactivation domain of CREB: a model for activator:coactivator interactions. Cell 91, 741–752.

Regier, J.L., Shen, F., and Triezenberg, S.J. (1993). Pattern of aromatic and hydrophobic amino acids critical for one of two subdomains of the VP16 transcriptional activator. Proc Natl Acad Sci U S A 90, 883–887.

Robinson, P.J., Trnka, M.J., Pellarin, R., Greenberg, C.H., Bushnell, D.A., Davis, R., Burlingame, A.L., Sali, A., and Kornberg, R.D. (2015). Molecular architecture of the yeast Mediator complex. Elife 4.

Saio, T., Guan, X., Rossi, P., Economou, A., and Kalodimos, C.G. (2014). Structural basis for protein antiaggregation activity of the trigger factor chaperone. Science 344, 1250494.

Schrodinger, LLC (2015). The PyMOL Molecular Graphics System, Version 1.8.

Schwieters, C.D., Kuszewski, J.J., Tjandra, N., and Clore, G.M. (2003). The Xplor-NIH NMR molecular structure determination package. J Magn Reson 160, 65–73.

Shen, Y., and Bax, A. (2013). Protein backbone and sidechain torsion angles predicted from NMR chemical shifts using artificial neural networks. J Biomol NMR 56, 227–241.

Sigler, P.B. (1988). Transcriptional activation. Acid blobs and negative noodles. Nature 333, 210–212.

Spitz, F., and Furlong, E.E. (2012). Transcription factors: from enhancer binding to developmental control. Nature reviews. Genetics 13, 613–626.

Swanson, M.J., Qiu, H., Sumibcay, L., Krueger, A., Kim, S.J., Natarajan, K., Yoon, S., and Hinnebusch, A.G. (2003). A multiplicity of coactivators is required by Gcn4p at individual promoters in vivo. Mol Cell Biol 23, 2800–2820.

Tamiola, K., and Mulder, F.A. (2012). Using NMR chemical shifts to calculate the propensity for structural order and disorder in proteins. Biochem Soc Trans 40, 1014–1020.

Tantos, A., Han, K.H., and Tompa, P. (2012). Intrinsic disorder in cell signaling and gene transcription. Mol Cell Endocrinol 348, 457–465.

Tian, Y., Schwieters, C.D., Opella, S.J., and Marassi, F.M. (2014). A practical implicit solvent potential for NMR structure calculation. J Magn Reson 243, 54–64.

Tompa, P., Davey, N.E., Gibson, T.J., and Babu, M.M. (2014). A million peptide motifs for the molecular biologist. Mol Cell 55, 161–169.

Tsai, K.L., Yu, X., Gopalan, S., Chao, T.C., Zhang, Y., Florens, L., Washburn, M.P., Murakami, K., Conaway, R.C., Conaway, J.W., et al. (2017). Mediator structure and rearrangements required for holoenzyme formation. Nature.

Unger, T., Nau, M.M., Segal, S., and Minna, J.D. (1992). p53: a transdominant regulator of transcription whose function is ablated by mutations occurring in human cancer. EMBO J 11, 1383–1390.

Walker, S., Greaves, R., and O’Hare, P. (1993). Transcriptional activation by the acidic domain of Vmw65 requires the integrity of the domain and involves additional determinants distinct from those necessary for TFIIB binding. Mol Cell Biol 13, 5233–3513.

Warfield, L., Tuttle, L.M., Pacheco, D., Klevit, R.E., and Hahn, S. (2014). A sequence-specific transcription activator motif and powerful synthetic variants that bind Mediator using a fuzzy protein interface. Proc Natl Acad Sci U S A 111, E3506–3513.

Waters, L., Yue, B., Veverka, V., Renshaw, P., Bramham, J., Matsuda, S., Frenkiel, T., Kelly, G., Muskett, F., Carr, M., et al. (2006). Structural diversity in p160/CREB-binding protein coactivator complexes. J Biol Chem 281, 14787–14795.

Weake, V.M., and Workman, J.L. (2010). Inducible gene expression: diverse regulatory mechanisms. Nature reviews. Genetics 11, 426–437.

Yang, B., Wu, Y.J., Zhu, M., Fan, S.B., Lin, J., Zhang, K., Li, S., Chi, H., Li, Y.X., Chen, H.F., et al. (2012). Identification of cross-linked peptides from complex samples. Nat Methods 9, 904–906.

Yang, F., Vought, B.W., Satterlee, J.S., Walker, A.K., Jim Sun, Z.Y., Watts, J.L., De Beaumont, R., Saito, R.M., Hyberts, S.G., Yang, S., et al. (2006). An ARC/Mediator subunit required for SREBP control of cholesterol and lipid homeostasis. Nature 442, 700–704.

Yoon, S., Qiu, H., Swanson, M.J., and Hinnebusch, A.G. (2003). Recruitment of SWI/SNF by Gcn4p does not require Snf2p or Gcn5p but depends strongly on SWI/SNF integrity, SRB mediator, and SAGA. Mol Cell Biol 23, 8829–8845.

Zor, T., De Guzman, R.N., Dyson, H.J., and Wright, P.E. (2004). Solution structure of the KIX domain of CBP bound to the transactivation domain of c-Myb. J Mol Biol 337, 521–534.

